# The role of Kinesin-1 in neuronal dense core vesicle transport and lifespan regulation in *C. elegans*

**DOI:** 10.1101/2024.03.25.586639

**Authors:** Anna Gavrilova, Astrid Boström, Nickolay Korabel, Sergei Fedotov, Gino B. Poulin, Victoria J. Allan

**Affiliations:** School of Biological Sciences, Faculty of Biology, Medicine and Health, University of Manchester, The Michael Smith Building, Rumford St, Manchester M13 9PT; Department of Mathematics, Faculty of Science and Engineering, The University of Manchester, Manchester M13 9PL, UK

**Author notes:** Corresponding author: Viki Allan.

**Keywords:** kinesin, dense core vesicle, neuron, *C. elegans*, lifespan

## Abstract

Fast axonal transport is crucial for neuronal function and is driven by kinesins and cytoplasmic dynein. We investigated the role of the kinesin-1 motor complex in dense core vesicle (DCV) transport in *C. elegans*, using mutants in kinesin light chains (*klc*-1 and *klc*-2) and the kinesin motor subunit (*unc-116*) expressing an *ida-1::gfp* transgene that labels DCVs in the ALA neuron. A reduced-function *unc-116(rf)* mutation greatly impaired DCV transport in both directions. A *klc-2(rf)* reduced-function mutation decreased DCV velocity in both directions and reduced the frequency of body bends during swimming. In contrast, the *klc-1(-)* null mutation had no effect on anterograde transport or swimming ability, but surprisingly it increased the speed of retrograde DCV transport. We also determined lifespan, finding that *klc-1(-)* or *klc-2(rf)* single mutants were wild-type whereas the *unc-116(rf)*, *ida-1::gfp* and *unc-116(rf)*; *ida-1::gfp* strains were short-lived. Strikingly, the *ida-1::gfp* transgenic synergistically interact with either *klc* mutant to extend lifespan compared to wild-type and parental strains. Our findings suggest that kinesin-1 not only influences anterograde and retrograde DCV transport but also plays a role in regulating lifespan.

## Introduction

Intracellular transport along microtubules drives the dynamic movement of organelles, vesicles and biomolecules over long distances within animal cells. It is crucial for maintaining cellular organization, homeostasis, and overall cell viability (Sleigh et al., 2019). This is particularly important in neurons since most proteins are synthesized and packaged in the cell body and must be effectively transported over substantial distances. A direct link has been observed between mutations in the microtubule motor proteins that drive this transport and a spectrum of nervous system disorders and neurological diseases, such as Alzheimer’s disease and Parkinson’s disease (Brady and Morfini, 2017; Hirokawa et al., 2010; Sleigh et al., 2019). Nonetheless, the relationship between disruptions in fast axonal transport and neuronal disorders is not fully understood.

In addition to relying on standard diffusion, intracellular cargos undergo active transport with the aid of two distinct ATP-dependent motor proteins: kinesins and dyneins (Guedes-Dias and Holzbaur, 2019; Hirokawa et al., 2010). Cytoplasmic dynein drives retrograde transport, towards microtubule minus ends, while the majority of kinesin family members support anterograde transport, towards microtubule plus ends (Hancock, 2014; Hirokawa et al., 2010). In axons, microtubules align parallel to the axon’s length, with the growing plus-ends extending away from the cell body towards the axon tip, while the minus-ends point towards the cell body (Baas and Lin, 2011). More than one kinesin type can transport the same cargo (e.g., Barkus et al., 2008; Grigoriev et al., 2007; Guardia et al., 2016; Gumy et al., 2017; Kulkarni et al., 2017; Laurent et al., 2018; Lim et al., 2017; Serra-Marques et al., 2020; Wiese et al., 2010; Zahavi et al., 2021); however, the specific contributions and roles of each motor, as well as the possibility of simultaneous engagement of different motor types in cargo transport, remain unclear for many cellular organelles.

Dense core vesicles (DCVs) are specialized membrane-bound organelles responsible for the transport of neuropeptides, monoamines, neurotrophic factors, hormones, insulin-related peptides, and preassembled components, to their sites of release in both axons and dendrites (Gondre-Lewis et al., 2012; Kuznetsov and Kuznetsov, 2017; Kwinter et al., 2009; Wong et al., 2012). This traffic is essential for neuronal growth, signalling, learning, development, movement and ageing (Gondre-Lewis et al., 2012; Hammarlund et al., 2008; Randi et al., 2023; Ripoll-Sanchez et al., 2023; Speese et al., 2007). DCVs are formed in the neuron soma, undergo fast axonal transport, and release their content in response to specific stimuli (Gondre-Lewis et al., 2012), with fusion occurring at both synaptic and non-synaptic sites (Bharat et al., 2017; Hammarlund et al., 2008; Schlager et al., 2010; Shakiryanova et al., 2006; Wong et al., 2012).

DCVs move in both directions in *C. elegans* (Edwards et al., 2015b; Goodwin and Juo, 2013; Goodwin et al., 2012; Laurent et al., 2018; Park et al., 2023; Zahn et al., 2004), *Drosophila* (Barkus et al., 2008; Lim et al., 2017; Shakiryanova et al., 2006; Sung and Lloyd, 2022) and rat (Arimura et al., 2009; Gumy et al., 2017; Lo et al., 2011; Schlager et al., 2010; Zahavi et al., 2021), with reversals being reported to occur primarily at axon terminals or synapses (Bharat et al., 2017; Guedes-Dias et al., 2019; Wong et al., 2012). Kinesin-3/UNC-104 has been shown to drive plus-end-directed DCV movement, and kinesin-3 mutations greatly reduce the number of DCVs entering the axon (Barkus et al., 2008; Edwards et al., 2015b; Goodwin and Juo, 2013; Goodwin et al., 2012; Gumy et al., 2017; Jacob and Kaplan, 2003; Laurent et al., 2018; Lo et al., 2011; Pack-Chung et al., 2007; Park et al., 2023; Sieburth et al., 2005; Zahn et al., 2004). Importantly, kinesin-1 is also a DCV motor in vertebrates and flies, leading to the hypothesis that kinesin-3 moves DCVs out of the cell body, then kinesin-1 takes over from, or works alongside, kinesin-3 for long distance movement (Arimura et al., 2009; Barkus et al., 2008; Gumy et al., 2017; Lim et al., 2017; Zahavi et al., 2021). In *C. elegans*, two peaks of anterograde velocity were seen for DCVs in ALA neurons (expressing IDA-1::GFP, a transmembrane phogrin homologue), with the faster being lost in an *unc-104* (kinesin-3) mutant (Zahn et al., 2004), again suggesting that another kinesin can drive DCV motility.

Here, we show that kinesin-1 is the second motor for this DCV transport in *C. elegans*. Kinesin-1 is a heterotetrameric molecular motor complex, consisting of two kinesin heavy chains (KHCs/KIF5) and two kinesin light chains (KLCs) (Anton et al., 2021; Cross and Dodding, 2019). In the genome of *C. elegans*, a single KHC is encoded by the *unc-116* gene, while two KLCs are encoded by the *klc-1* and *klc-2* genes (Sakamoto et al., 2005). KLCs play a crucial role in modulating motor binding to cargo and are actively involved in activating the motor (Cross and Dodding, 2019; Hirokawa et al., 2010). To test the roles of the two distinct KLCs and KHC in DCV transport, we analysed the movement of IDA-1::GFP-labelled DCVs in three strains bearing mutations in the *klc-1*, *klc-2* and *unc-116* (KHC) genes. We find that DCV movement in both directions is severely compromised by the *unc-116* mutation, and partially affected by the *klc-2* mutation. Surprisingly, dynein-driven DCV motility is faster in the *klc-1* mutant than in the wild-type background, while plus-end-directed movement is unaffected. While the *klc-1* and *-2* mutant worms had normal longevity, the *unc-116* mutant worms were short-lived. Unexpectedly, the expression of the *ida-1::gfp* transgene also reduced worm lifespan, but combining *ida-1::gfp* with either *klc* mutation gave animals with a longer lifespan than wildtype worms or either parental strain. This highlights a novel role for kinesin-1 in the control of longevity in certain genetic backgrounds.

## Results

The IDA-1 (related to *i*slet cell *d*iabetes *a*utoantigen) protein is the *C. elegans* orthologue of mammalian type-1 diabetes auto-antigen proteins IA-2 (Insulinoma Associated protein-2) and phogrin (IA-2*β*) (Cai et al., 2001; Zahn et al., 2001). Expression of IDA-1::GFP from its endogenous promoter revealed localisation to DCVs in the ALA, VC, HSN and PHC neurons, and the uv1 neurosecretory cell (Cai et al., 2004; Zahn et al., 2004), although RNAseq and *ida-1*-promoter-driven GFP expression suggest *ida-1* is expressed more widely at lower levels (Cai et al., 2001; Cao et al., 2017; Taylor et al., 2021; Zahn et al., 2001). Here, we used a strain expressing IDA-1::GFP from two integrated extrachromosomal arrays under the control of the *ida-1* promoter (giving ∼2-3 fold over-expression of IDA-1 protein), which gives strong GFP labelling of DCVs in the ALA neuron (Zahn et al., 2004), as shown in Figure 1A. The ALA neuron is ideal for imaging, as it extends two axons down each side of the worm from the cell body in the head. During development, the ALA axons guide the extension of the primary dendrites of the PVD neuron and interact closely with the CAN neuron to form the lateral nerve tract that runs alongside the excretory canal (Ramirez-Suarez et al., 2019). It has relatively few chemical synapses, but instead both receives and transmits signals via neuropeptides (Iannacone et al., 2017; Konietzka et al., 2020; Nath et al., 2016; Nelson et al., 2014; Randi et al., 2023; Ripoll-Sanchez et al., 2023).

**Figure 1.**
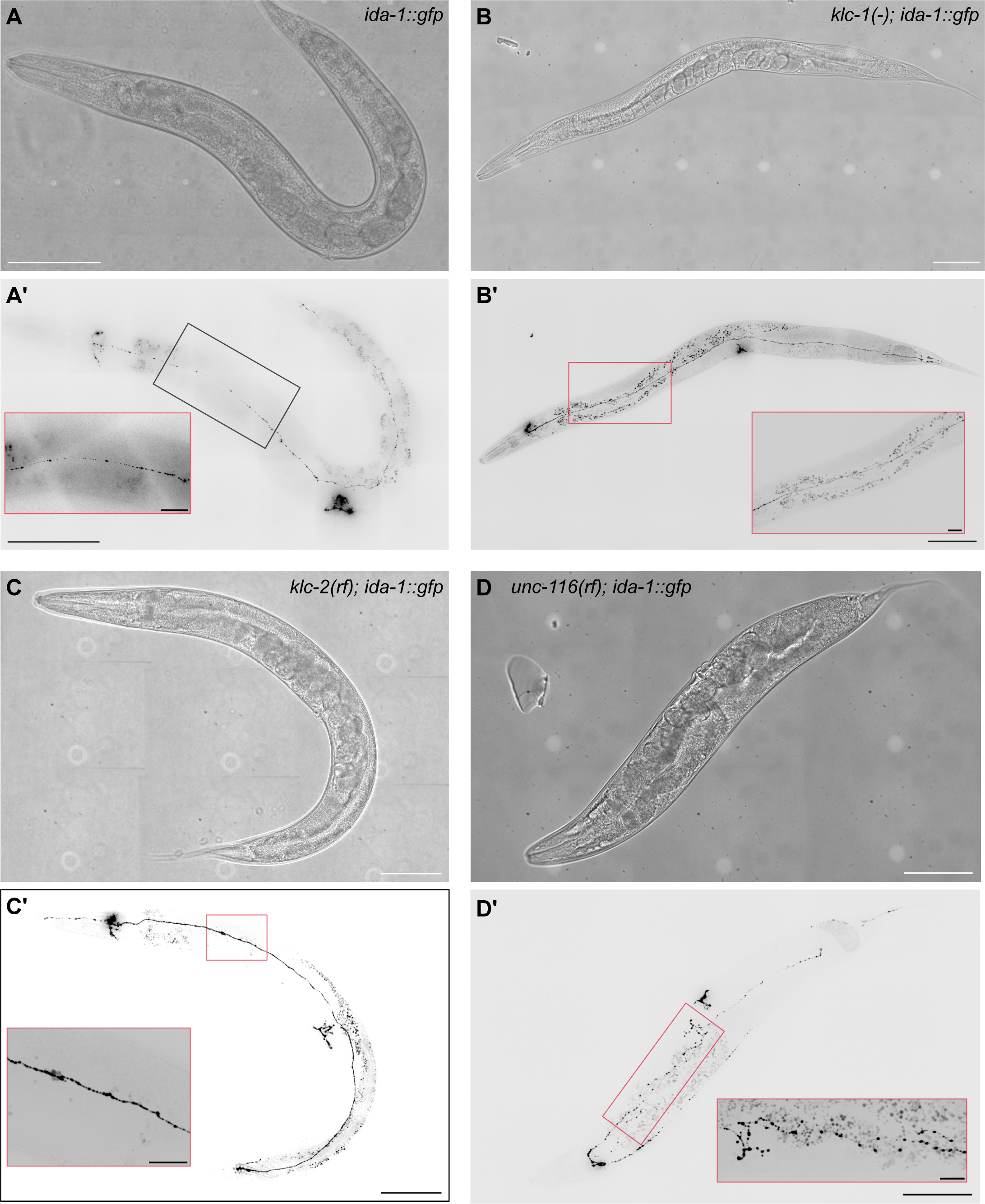
Worm morphology and dense core vesicle (DCV) distribution. Spinning disk brightfield and fluorescence images of an A) *ida-1::gfp*, B) *klc-1(-); ida-1::gfp*, C) *klc-2(rf); ida-1::gfp*, and D) *unc-116(rf); ida-1::gfp* worms showing the DCVs (inverted LUT). Scale bars are 100 µm. An enlarged section of each ALA neuron is shown in the red boxes. The lighter spots in the brightfield images are artefacts generated by the tiling of multiple images. Scale bars = 20 µm. The larger particles visible outside the ALA neuron are auto-fluorescent granules, mainly located in the gut. Images were collected using a 20 X objective for *ida-1::gfp* and a 40 X objective for the other strains.

### Alterations in ALA neuron morphology in an *unc-116* mutant

To test the role of kinesin-1 in DCV motility in *C. elegans*, the *ida-1::gfp* strain was crossed with kinesin-1 reduced-function mutants in kinesin heavy chain (*unc-116(rh24sb79)*: referred to as *unc-116(rf)* herein) (Patel et al., 1993; Yang et al., 2005), kinesin light chain 2 (*klc-2(km11)*: referred to here as *klc-2(rf)*) (Sakamoto et al., 2005) and a kinesin light chain 1 deletion mutant (*klc-1(ok2609)*: referred to as *klc-1(-)*) (Consortium, 2012). Imaging IDA-1::GFP in these new strains revealed normal ALA morphology and DCV distribution in the *klc-1(-)* and *klc-2(rf)* mutant strains (Fig. 1A-C). In contrast, *unc-116(rf); ida-1::gfp* worms often displayed branching of the ALA axons (Figs. 1D, S1). It is known that *unc-116* mutant strains can have alterations in dendrite branching in the PVD neuron that are driven by changes in microtubule orientation (Aguirre-Chen et al., 2011; Harterink et al., 2018; Taylor et al., 2015). In contrast, microtubule organisation in axons was normal in *unc-116* mutants in all axons tested (Harterink et al., 2018; Yan et al., 2013). Inappropriate branching of axons has been observed in zebrafish (Auer et al., 2015), but it has not been reported in *C. elegans* kinesin-1 mutants to our knowledge.

### Effects of kinesin-1 mutants on DCV motility

To characterise the motile properties of DCVs in the ALA neuron of these strains, we imaged single z-planes by spinning disk confocal microscopy. From each video, the position of DCVs over time was displayed as kymograph of the fluorescence signal along the axon (Figs. 2 and S2). Consistent with a previous report (Zahn et al., 2004), in the wild-type background we observed DCVs moving in both directions, but rarely changing direction (Fig. 2A, S2A; Movie 1). In addition, there were notable spots where particles clustered together and did not move, which likely correspond to the periodic swellings observed by EM along the length of the ALA neuron that contain clusters of DCV, some of which may occur at *en passant* synapses (Ramirez-Suarez et al., 2019). When approaching such a cluster, the moving DCV could either stop or pass through. Clusters were sometimes seen to shed a DCV that then moved off in either direction.

**Figure 2.**
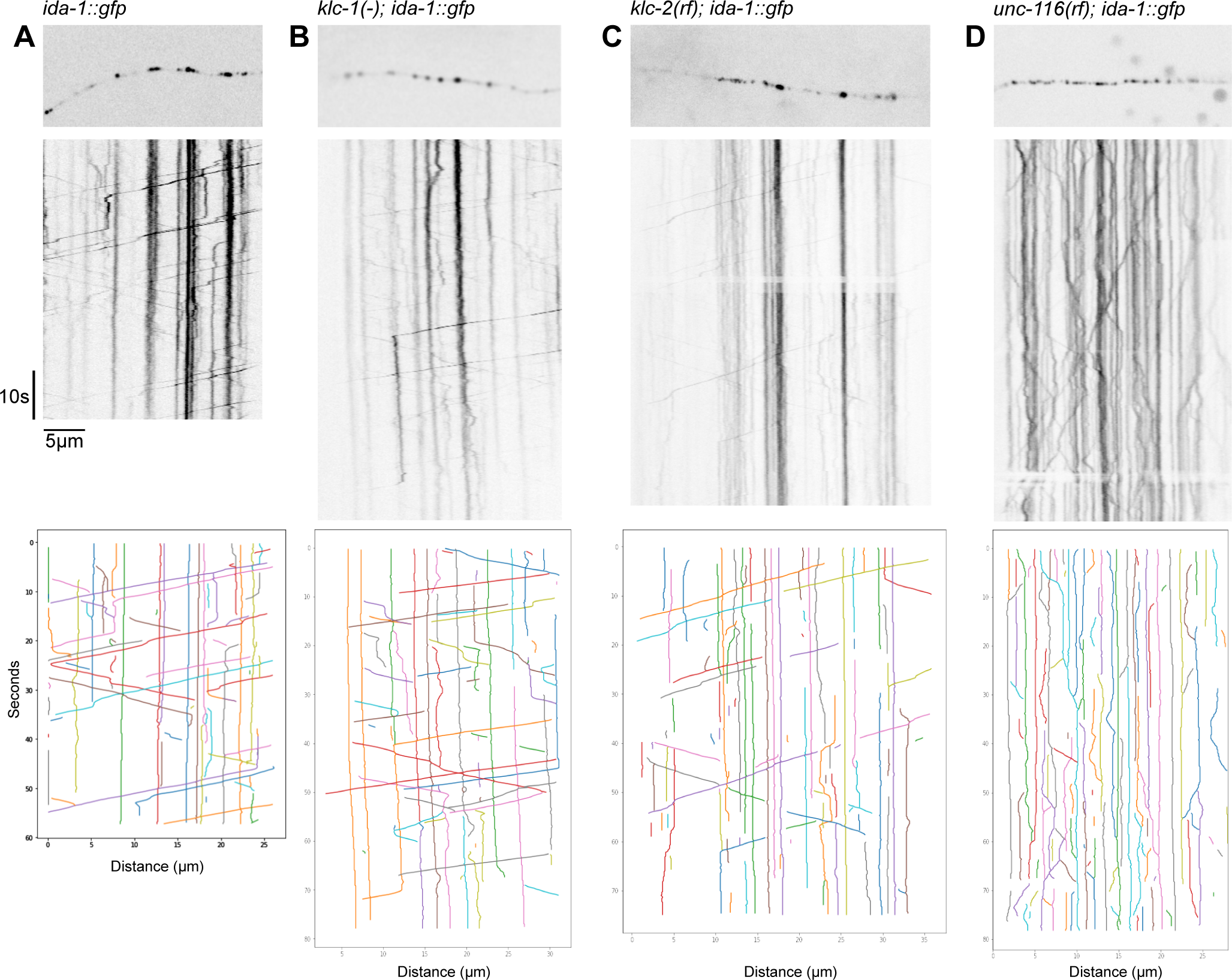
Kymographs of DCV movement in the ALA neuron. The initial frame of each movie is shown on top, with the kymograph below. A) *ida-1::gfp*, B) *klc-1(-); ida-1::gfp*, C) *klc-2(rf); ida-1::gfp*, and D) *ida-1::gfp*; *unc-116(rf)*. The nerve terminal (microtubule plus ends) is on the right. Kinesin-driven DCVs correspond to lines sloping from top left to bottom right, with dynein-driven lines sloping from top right to bottom left. Vertical lines indicate stationary DCVs (individual or clusters). Scale bars = 5 µm.

To obtain velocities of DCV movement, kymographs were automatically tracked using KymoButler (Jakobs et al., 2019) then analysed using custom-written Python scripts. A plot of DCV displacement vs time for all tracks (Fig. 3A) revealed that dynein-driven motion was highly processive (DCVs moving without stopping), with an average velocity of 2.14 ± 0.77 µm/s (Table 1). Plus-end-directed movement was less processive and had an average velocity of 1.4 ± 0.68 µm/s (Table 1). The distribution of segment velocities (Fig. 4A) broadly resembles that reported previously for a small number of DCVs in the ALA neuron (Zahn et al., 2004). Gaussian fitting revealed three peaks (Fig. S5), which likely correspond to kinesin-1 (slow), kinesin-3/UNC-104 (fast) and a mixture of the two (intermediate), as has been previously observed for mixtures of kinesin-1 and -3 motors, both in vitro and in DCV movement (Arpag et al., 2019; Arpag et al., 2014; Gumy et al., 2017; Norris et al., 2014; Serra-Marques et al., 2020; Soppina et al., 2014). Scoring the tracks according to their behaviour (plus- or minus-end-directed only, bidirectional, stationary, or combinations thereof, as summarised in Fig. S3) revealed that 21.5% of DCVs moved without pausing, 11.9% moved with pauses, and 66.6% of DCVs were stationary for the duration of the movie. In total, 12.2% of DCVs moved for some or all the time towards the plus ends, with 22.7% having dynein-driven runs (Fig. S3A).

**Figure 3.**
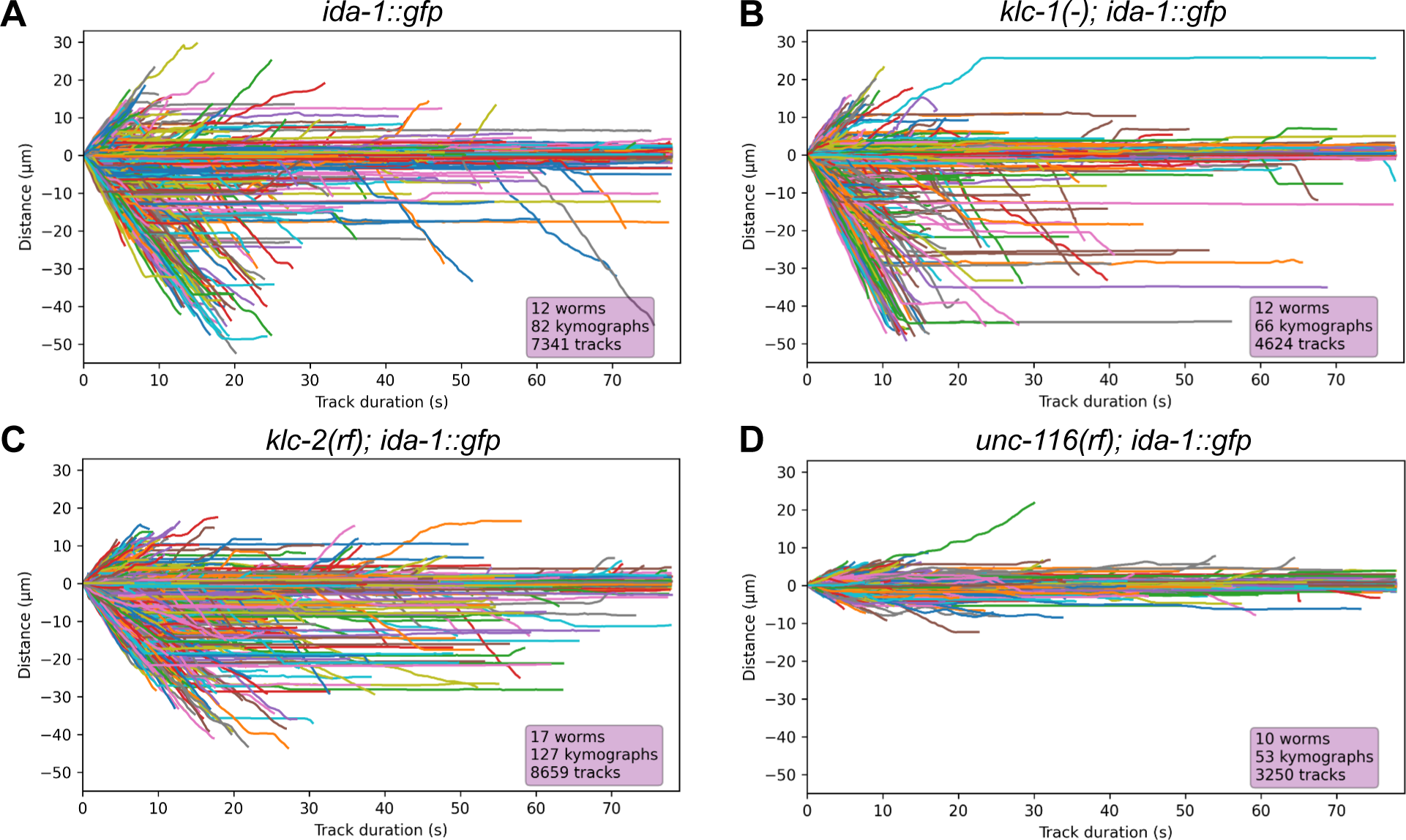
Visualisation of individual DCV tracks in the ALA neuron. Displacement plots (distance moved vs time) showing the movement of DCVs in the ALA neuron in: A) *ida-1::gfp*, B) *klc-1(-); ida-1::gfp*, C) *klc-2(rf); ida-1::gfp*, and D) *unc-116(rf); ida-1::gfp*.

**Figure 4.**
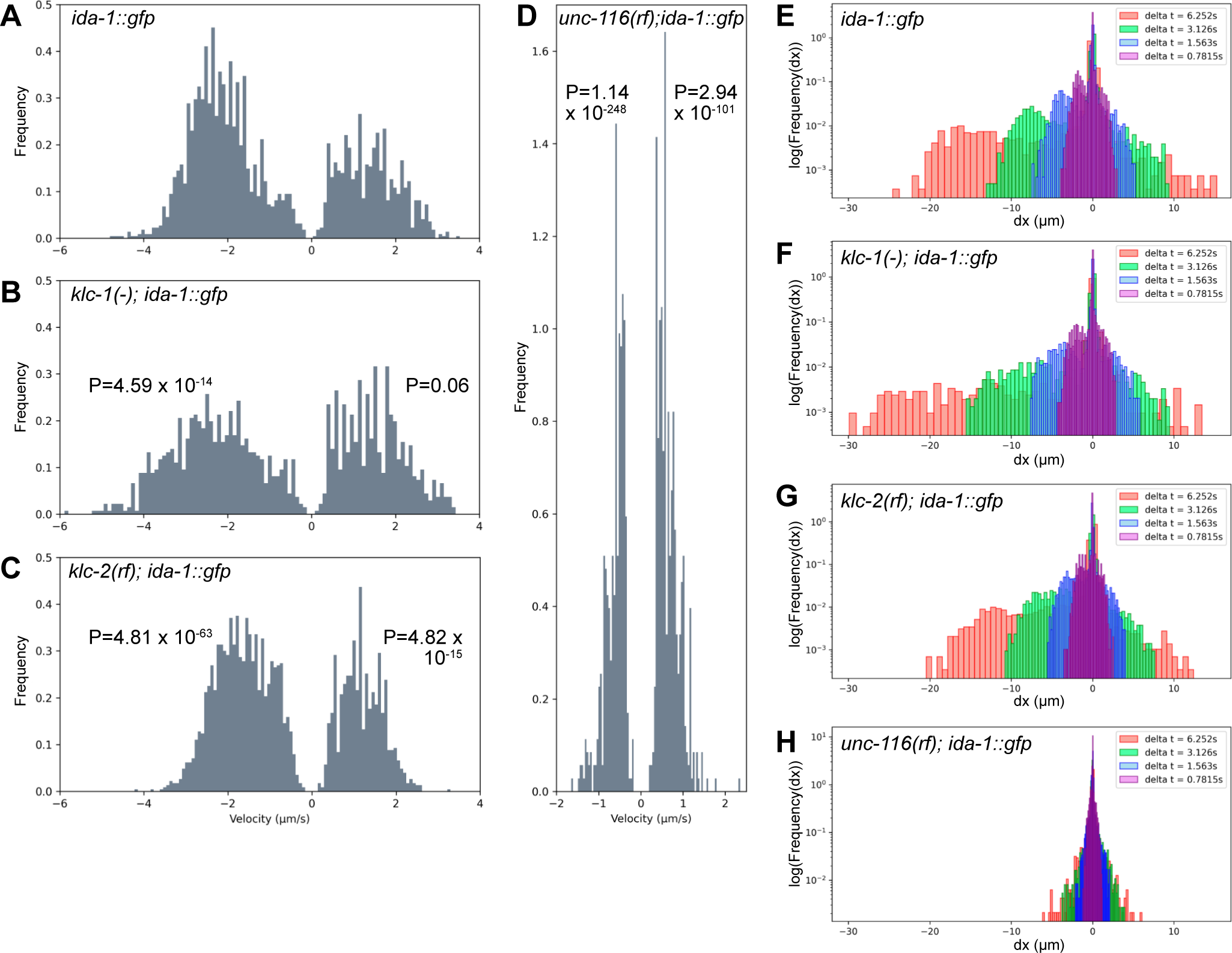
Velocity and displacement distribution plots of DCV movement. The distribution of velocities of active segments are plotted for A) *ida-1::gfp*, B) *klc-1(-); ida-1::gfp*, C) *klc-2(rf); ida-1::gfp*, and D) *unc-116(rf); ida-1::gfp*. The probability density functions (Frequency) were estimated from histograms with a bin size of 100; negative velocities are for dynein-driven movement; positive velocities are for movement towards the tail (microtubule plus ends). Note that the Y-axis scale is the same in A-D. The number of segments used to generate velocities is given in Table 1. P-values from pairwise Kolmogorov-Smirnov tests for velocity distribution of DCVs in the different strains are shown for both directions of movement (see also Table S1). This data is replotted as the distribution of displacements (on a log scale) within 5, 10, 20 or 40 frames (E-H).

**Table 1.** Summary of DCV motility analysis. Track segments with active movement in the indicated direction were used to generate the velocities (plotted in Fig. 4) summarised here.

|  | *ida-1::gfp* | *klc-1(-);*<br>*ida-1::gfp* | *klc-2(rf);*<br>*ida-1::gfp* | *unc-116 (rf);*<br>*ida-1::gfp* |
| --- | --- | --- | --- | --- |
| Number of worms;<br>kymographs | 12; 82 | 12; 66 | 17; 127 | 10; 53 |
| Number of tracks | 7341 | 4624 | 8659 | 3250 |
| Number of active<br>retrograde segments | 1719 | 816 | 1913 | 398 |
| Mean retrograde<br>velocity ($\mu\text{m/s} \pm \text{SD}$) | 2.14 ($\pm 0.77$) | 2.38 ($\pm 1.03$) | 1.66 ($\pm 0.67$) | 0.63 ($\pm 0.24$) |
| Median retrograde<br>velocity ($\mu\text{m/s}$) | 2.19 | 2.35 | 1.66 | 0.59 |
| Maximum retrograde<br>velocity ($\mu\text{m/s}$) | 4.81 | 5.89 | 4.22 | 1.65 |
| Number of active<br>anterograde segments | 940 | 643 | 1100 | 485 |
| Mean anterograde<br>velocity ($\mu\text{m/s} \pm \text{SD}$) | 1.4 ($\pm 0.68$) | 1.53 ($\pm 0.73$) | 1.2 ($\pm 0.48$) | 0.67 ($\pm 0.26$) |
| Median anterograde<br>velocity ($\mu\text{m/s}$) | 1.37 | 1.5 | 1.18 | 0.59 |
| Maximum anterograde<br>velocity ($\mu\text{m/s}$) | 3.53 | 3.43 | 3.3 | 2.35 |

The motile behaviour of DCVs in the *klc-1(-)* and *klc-2(rf)* mutants (Fig. 2B,C, S2B,C; Movies 2 and 3 respectively) was broadly similar to wild-type. However, in the *klc-2(rf)* mutant the slopes of the kymograph tracks in both directions were clearly different to those in the wild-type background (Fig. 3C). Analysis revealed that the mean DCV velocity was reduced in both directions (1.2 ± 0.48 µm/s plus-end-directed; 1.66 ± 0.67 µm/s dynein-driven: Table 1). Comparison of the distribution of the individual segment velocities showed that both directions were significantly different to those in wild-type *ida-1::gfp* worms (Fig. 4C; Table S1), and gaussian fitting revealed the loss of the fastest plus-end-directed segments (Fig. S5). The percentage of DCVs moving in each direction and those remaining stationary were similar (±1.5%) to wild-type, however (Fig. S3C).

In contrast to the *klc-2(rf)* mutant, the complete loss of *klc-1* had only a small, statistically insignificant effect on plus-end-directed mean velocity (Figs. 3B, 4B; Tables 1,S1) and the distribution of segment velocities (Fig. S5). Surprisingly, dynein-driven DCVs moved significantly *faster* in the *klc-1(-)* mutant than wild-type, with an average velocity of 2.38 ± 1.03 µm/s (Table 1). The velocity distribution plot (Fig. 4B) revealed an increase in the proportion of dyein-driven DCVs moving at very high speeds, and the maximum retrograde DCV velocity also increased from 4.81 to 5.89 µm/s (Table 1). Again, the percentages of DCVs moving in each direction or stationary were only slightly different (±4.4%) to those in the wild-type and *klc-2(rf)* (Fig. S3). As an alternative way of demonstrating the difference in behaviour of the fastest-moving DCVs, we replotted the data to show the number of DCVs that reached a given displacement after 5, 10, 20 or 40 frames (Fig. 4E-G). This clearly shows the greater distance travelled by dynein-driven DCVs within a set time in the *klc-1(-)* mutant compared to the wild-type and *klc-2(rf)* mutant.

A much greater effect on DCV motility was seen in the *unc-116(rf)* mutant background. Although movement still occurred in both directions, the kymographs revealed hesitant, slower motion (Figs. 2D, 3D, S2D; Movie 4). Analysis of the automatically segmented kymographs confirmed a significant reduction in mean velocity, to 0.67 ± 0.26 µm/s for plus-end-directed motion and 0.63 ± 0.24 µm/s for dynein-driven motility (Tables 1, S1). Likewise, the velocity distribution plot showed a much narrower range of values (Fig. 4D: note that the Y axis scale is the same for graphs A-D), with a maximum of +2.35 or -1.65 µm/s (Table 1). Furthermore, the displacements were shorter (Fig. 3D, 4H) and there was an increase in the percentage of stationary DCVs with a corresponding decrease in the percentage of both plus- and minus-end-directed DCV movement (Fig. S3D). Gaussian fitting of the segment velocities showed very different distributions, with new, slow peaks in both directions (Fig. S5)

Since kinesin-1 is a plus-end-directed motor that would transport DCVs from the cell body towards the axon tip, we wondered if DCV motility changed with increasing distance from the cell body. We therefore separated the data into 4 groups according to where in the ALA axon it was collected, then plotted the displacement vs time for each track (Fig. S4). Since the *unc-116(rf)* and *klc-2(rf)* worms have a shorter body length (1.12 mm for *ida-1::gfp*; 1.12 mm for *klc-1(-); ida-1::gfp*; 0.99 mm for *klc-2(rf); ida-1::gfp*, and 0.77 mm for *unc-116(rf); ida-1::gfp*, n = 100), there were no kymographs collected more than 622 or 696 µm from the ALA cell body in *unc-116(rf)* and *klc-2(rf)* worms, respectively (Fig. S4C,D). This analysis showed that DCV behaviour was not affected by position along the axon.

### Effect of kinesin-1 mutations on *C. elegans* swimming

Since different kinesin-1 mutants had distinct effects on DCV motility, indicating neuronal dysfunction, we decided to test whether the worms’ ability to move is compromised. In addition, swimming ability of *ida-1* null worms is reduced (Fatima et al., 2014), raising the question of what happens when IDA-1 protein levels are increased by the presence of the *ida-1::gfp* transgene. We assessed mobility by performing swimming (also called thrashing) assays and calculated the number of body bends per second that adult worms have when swimming (Fig. 5, Table S2). The bending frequency of *ida-1::gfp* worms was the same as wildtype N2 worms, as was that of the *klc-1(-)* mutant with or without expression of *ida-1::gfp*. As expected given their uncoordinated phenotype, the bending frequency of both *unc-116(rf)* and *unc-116(rf); ida-1::gfp* was extremely low and not significantly different. The *klc-2(rf)* mutant on its own had a lower rate of body bends than N2, consistent with their slightly *unc* phenotype when crawling (Sakamoto et al., 2005). Surprisingly, however, this was improved significantly (Table S1) by the presence of the *ida-1::gfp* transgene. The *klc-1(-); ida-1::gfp* double mutant had no effect on body bends, presumably because single mutant *klc-1(-)* has bending frequency same as N2 and therefore could not be improved upon. Thus, kinesin-1 containing KLC-2 is required for worm motility, which is an indicator of longevity (Hahm et al., 2015; Oswal et al., 2022).

**Figure 5.**
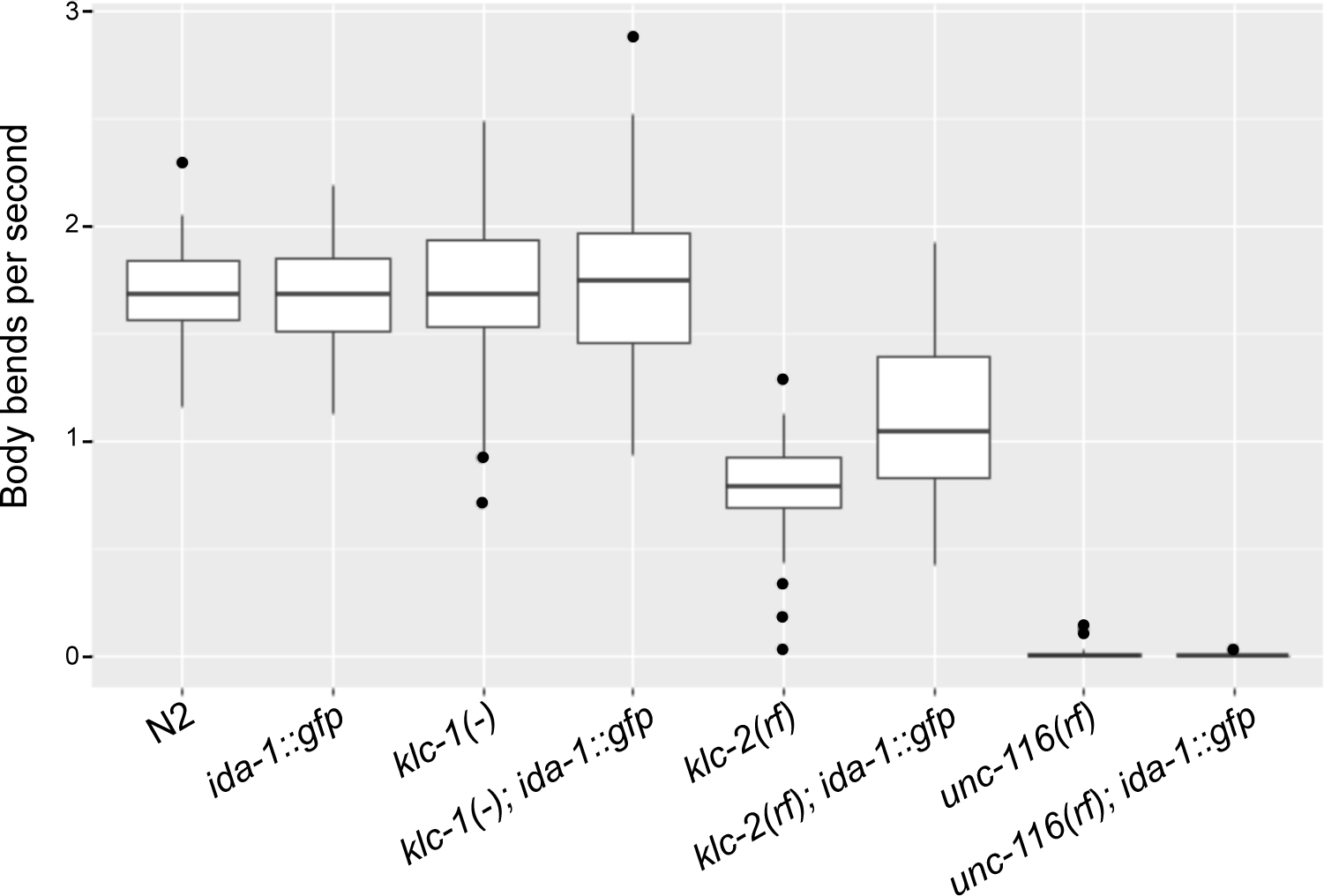
Effect of kinesin-1 mutations and *ida-1::gfp* expression on the mobility of *C. elegans*. A) Body bends per second of N2, *ida-1::gfp*, *klc-1(-)*, *klc-2(rf)*, *unc-116(rf)* and *klc-1(-)*, *klc-2(rf)*, or *unc-116(rf)* strains expressing *ida-1::gfp*. Worms were analysed as day 1 adults in M9 buffer. The boxplot displays the median and two hinges, corresponding to the first and third quartiles. The whiskers extend from the hinge to the largest or smallest value within 1.5 times the interquartile range (IQR) from the hinge. All “outlying” points are plotted individually. Statistical analysis is provided in Table S2.

### Effect of kinesin-1 mutations and expression of the *ida-1::gfp* transgene on lifespan

The genetic interaction between the *klc-2(rf)* mutation and the ida-1::gfp transgene that led to enhanced worm motility indicates improved health, which could translate into an increased lifespan (Hahm et al., 2015; Oswal et al., 2022). We therefore directly assessed whether lifespan could be improved in any of the kinesin-1 mutants by the presence of *ida-1::gfp* (Fig. 6; Table S3).

**Figure 6.**
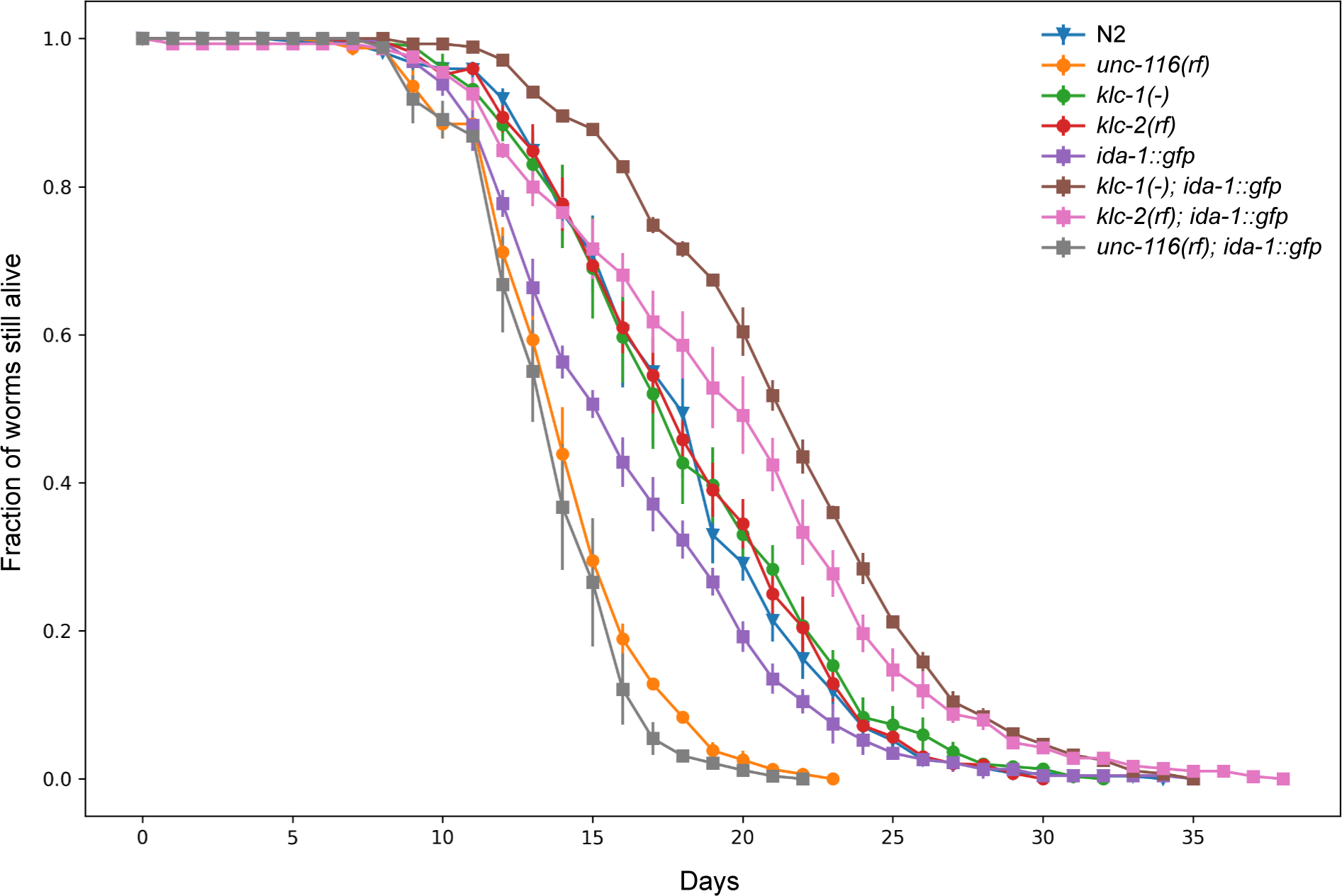
Effect of kinesin-1 mutations and *ida-1::gfp* expression on the lifespan of *C. elegans*. Survival plots showing the percentage of live worms as a function of time for N2, *ida-1::gfp*, *klc-1(-)*, *klc-2(rf)*, *unc-116(rf)* and the *klc-1(-)*, *klc-2(rf)*, or *unc-116(rf)* strains expressing *ida-1::gfp*. Day 0 is the L4 stage of the worm. Each assay started with 100-130 worms of each strain and was repeated three times. Bars represent standard error. The online application for survival analysis, OASIS 2 (Han et al., 2016), was used for analysis. A Log Rank Test was used to compare pairs of survival functions (Table S3).

The wild-type N2, *klc-1(-)* and *klc-2(rf)* worms had similar lifespans (50% survival at day 18 for all strains) meaning that these kinesin light chain mutations do not affect longevity. Interestingly, the lifespan of *ida-1::gfp* worms (50% survival at 16 days) was shorter than N2 suggesting that expression of this transgene has an important negative impact on lifespan. Both *unc-116(rf)* and the *unc-116(rf)*; *ida-1::gfp* strains had greatly reduced lifespans (50% survival at 14 days) compared to N2 and *ida-1::gfp*, with the *unc-116(rf); ida-1::gfp* being somewhat worse than *unc-116(rf)* alone at later times (Fig. 6; Table S3). Striking, however, expression of the *ida-1::gfp* transgene in either *klc* mutant gave a significant increase in lifespans (50% survival at 22 and 20 days, respectively) compared to N2 or the *klc* single mutants. Notably, these strains are dramatically longer lived than the *ida-1::gfp* parental strain, with the biggest effect seen in the *klc-1(-)* mutant background. Therefore, *klc-1(-)* and *klc-2(rf)* mutants genetically interact with the short-lived *ida-1::gfp* transgene strain, to synergistically increase lifespan when compared to their parental strains.

## Discussion

### Kinesin-1 is a motor for DCV movement

Axonal transport of large organelles such as DCVs through narrow calibre axons is a challenging process (Maday et al., 2014; Sabharwal and Koushika, 2019). It is well established that kinesin-3 motors—UNC-104 in worms and flies, and KIF1 in vertebrates— are key drivers of this movement, since DCVs are mainly stuck in the neuron cell body when its function is disrupted (Barkus et al., 2008; Edwards et al., 2015b; Goodwin and Juo, 2013; Goodwin et al., 2012; Gumy et al., 2017; Laurent et al., 2018; Park et al., 2023; Zahn et al., 2004). By studying DCV transport in the *C. elegans* ALA neuron we show that kinesin-1 is also vital, because DCVs move much more slowly and hesitantly in worms with a reduced-function mutation in the motor subunit of kinesin-1, UNC-116. Nevertheless, DCVs in the *unc-116(rf)* worms do enter the axon, are evenly distributed along its length, and still move in both directions to a degree. In contrast, most DCVs remained in the ALA cell body in an *unc-104* mutant, whilst those that entered the axon moved well, albeit more slowly (Zahn et al., 2004). In addition, two organelles known to be cargoes for just kinesin-1, mitochondria and calsyntenin-positive vesicles, remain in the neuronal cell body when kinesin-1 function is disrupted (Araki et al., 2007; Konecna et al., 2006; Pilling et al., 2006; Rawson et al., 2014; Sure et al., 2018; Zhao et al., 2021). Altogether, these data strongly suggests that kinesins-1 and -3 work in tandem to drive DCV along the ALA neuron, as has been observed in rodent and *Drosophila* neurons (Barkus et al., 2008; Gumy et al., 2017; Lim et al., 2017; Zahavi et al., 2021). *C. elegans* DCVs are therefore another example of a cargo whose transport involves more than one kinesin type (e.g., Barkus et al., 2008; Grigoriev et al., 2007; Guardia et al., 2016; Gumy et al., 2017; Kulkarni et al., 2017; Laurent et al., 2018; Lim et al., 2017; Prevo et al., 2015; Serra-Marques et al., 2020; Wiese et al., 2010; Zahavi et al., 2021). Combining kinesins with different mechanochemical properties is thought to provide optimal cargo transport in complex cellular environments (Arpag et al., 2019; Gumy et al., 2017; Lim et al., 2017; Prevo et al., 2015).

The motility of DCVs in wildtype animals revealed a wide spread of velocities that were fitted by three Gaussian distributions (Fig. S5). Similar distributions are seen for DCV movement in mouse and rat neurons (Gumy et al., 2017) and MRC5 cells (Gumy et al., 2017; Serra-Marques et al., 2020). This fits with kinesin-1 being a slow motor whereas kinesin-3 motors are fast (Arpag et al., 2019; Arpag et al., 2014; Glomb et al., 2023; Guedes-Dias et al., 2019; Gumy et al., 2017; Norris et al., 2014; Serra-Marques et al., 2020; Soppina et al., 2014), and implies that many DCV runs are driven by one motor or the other. The presence of a population with intermediate velocity could reflect engagement of both motors at once, or rapid switching between them, as previously suggested (Serra-Marques et al., 2020). Notably, however, the plus-end-directed velocities in the *unc-116(rf)* mutant, which still has wildtype UNC-104 motors, is drastically slowed to a velocity not seen in normal axons, despite UNC-104 being a fast motor. This may be due to kinesin-3 motors detaching more readily from microtubules than kinesin-1 (Arpag et al., 2019; Arpag et al., 2014; Norris et al., 2014), particularly under load. The presence of the partially functional UNC-116 motor might also act as a brake, making detachment more likely. Alternatively, since dimerization of kinesin-3 motors has been shown to make them super-processive (Soppina et al., 2014), perhaps UNC-104 is unable to dimerise in the absence of functional UNC-116. Another possibility there is not enough UNC-104 present on the DCVs to be able to drive their rapid movement through the highly crowded axonal environment without the help of kinesin-1, as has been observed for DCV motility in MRC5 cells (Serra-Marques et al., 2020).

### Effect of *unc-116* and *klc-2* mutants on dynein-driven DCV motility

A striking feature of DCV motility in the ALA neuron is its bidirectional nature, which fits with previous reports for DCVs in a range of organisms (Barkus et al., 2008; Edwards et al., 2015b; Goodwin and Juo, 2013; Goodwin et al., 2012; Gumy et al., 2017; Laurent et al., 2018; Lim et al., 2017; Park et al., 2023; Schlager et al., 2010; Sung and Lloyd, 2022; Zahn et al., 2004). DCVs in the ALA axon switched direction infrequently: therefore, in our observation period each DCV was more likely to continue in one direction, sometimes with pauses. Indeed, reversals of direction and pauses have been observed mainly at *en passant* boutons and axon terminals (Bharat et al., 2017; Guedes-Dias et al., 2019; Wong et al., 2012).

We find that the reduced-function mutants of *unc-116* and *klc-2* decreased the velocity and processivity of both directions of movement, despite the presence of wild-type UNC-104. This effect was strongest in the *unc-116(rf)* mutant, where mean velocities dropped to 0.63 µm/s and 0.67 µm/s, a drop of 71% for dynein and 52% for plus-end-directed motion, respectively. This effect, termed “the paradox of co-dependence” (Hancock, 2014), has been seen in many different situations, ranging from acute antibody-mediated inhibition of one motor (e.g. Brady et al., 1990; Uchida et al., 2009; Yi et al., 2011) through to function-modifying mutations, deletions or depletion (e.g., Ally et al., 2009; Encalada et al., 2011; Iacobucci et al., 2014; Ling et al., 2004 and see below). For example, kinesin heavy chain disruptions caused an increased number of axonal swellings and caused organelle jams that affected both anterograde and retrograde transport in *Drosophila* (Haghnia et al., 2007; Hurd and Saxton, 1996; Lim et al., 2017; Neisch et al., 2017; Pilling et al., 2006). This interrelationship has also been reported for many cargos in *C. elegans* (e.g., Celestino et al., 2022; Edwards et al., 2015a; Hoerndli et al., 2013; Hoerndli et al., 2015), including DCVs in the PQR sensory neuron (Laurent et al., 2018) as well as in the ALA neuron, as described here.

It is common to see similar reductions in velocity for both directions following kinesin-1 disruption, not just the number of movements (e.g. Celestino et al., 2022; Ding et al., 2022; Gumy et al., 2017; Hoerndli et al., 2013). In contrast, the rate of dynein-driven DCV movement in the ALA neuron was normal in an *unc-104* mutant despite the loss of the fastest plus-end-directed motion (Zahn et al., 2004). Normal dynein-driven rates have been seen in other situations where kinesin-3 function is impaired (Laurent et al., 2018; Ou et al., 2010). This may fit with the observation that a kinesin-3 (KIF13B) on DCVs does not engage in a tug-of-war with dynein, whereas kinesin-1 does (Serra-Marques et al., 2020). However, other studies have reported reduced dynein DCV speeds in kinesin-3 mutants (Barkus et al., 2008; Lo et al., 2011). These differing results may be due to the different motor mutations or inhibition methods used and cargoes analysed.

### Loss of KLC-1 leads to faster dynein movement of DCVs

A striking finding in this study is that whilst the null *klc-1* mutation did not affect the anterograde movement of DCVs, it actually increased the speed of retrograde transport. The *klc-1* gene is expressed at very low levels in the ALA neuron (Taylor et al., 2021) so it may play only a minor role in DCV transport, if any. How might its loss enhance dynein-driven DCV motility specifically?

Physical crowding within neurons is an important constraint on cargo motility (Maday et al., 2014; Sabharwal and Koushika, 2019). One possibility is that absence of KLC-1 may change the physical environment in the ALA neuron in a way that facilitates dynein processivity indirectly. The limited width of axons means that moving DCVs occupy most of that diameter. For example, published ALA EM images (Ramirez-Suarez et al., 2019) reveal axon diameters of 0.087-0.54 µm whilst that of DCVs is ∼0.074 µm. In addition, stationary DCV accumulations are seen all along the axon in wild-type neurons (Figs. 1,2,S1,S2; (Bharat et al., 2017; Hammarlund et al., 2008; Schlager et al., 2010; Shakiryanova et al., 2006; Wong et al., 2012; Zahavi et al., 2021), which many DCVs move straight through whereas others pause temporarily or stop there. Such clusters may correspond to *en passant* synapses or non-synaptic release sites (Bharat et al., 2017; Hammarlund et al., 2008; Schlager et al., 2010; Wong et al., 2012), or be temporary traffic jams at regions with accumulations of various membrane cargoes (Iacobucci et al., 2014; Sood et al., 2018). They may also occur at microtubule ends, where motors and cargo may pause (Guedes-Dias et al., 2019; Yogev et al., 2016). The detachment of motors from microtubule ends can be a very slow process, leading to the accumulation of motors and, consequently, the formation of bottlenecks and jams (Leduc et al., 2012). Furthermore, stationary cargo accumulations in neurons often occur in actin-rich regions (Sood et al., 2018), and depolymerisation of actin increased DCV speeds in mouse cultured neurons, although directionality was not assessed (Bharat et al., 2017). Interestingly, mutation in the *C. elegans* neurofilament protein TAG-63 increased the retrograde, but not anterograde, speed of vesicles carrying synaptobrevin (Bhan et al., 2020). Importantly, neurofilaments have been shown to be cargoes of kinesin-1 and dynein in mouse (Uchida et al., 2009).

Why, though, would changing the physical environment favour dynein over plus-end-directed DCV transport? One possibility is that the latter may be driven by both two motors with distinct biophysical properties—UNC-104 and UNC-116—which may give more scope for dealing with obstructions (reviewed in (Maday et al., 2014; Sabharwal and Koushika, 2019)). Another possibility is that neuronal activity is altered in some way in the *klc-1(-)* mutant either to promote dynein activation or reduce the likelihood of dynein-driven DCVs pausing. Notably, in *Drosophila*, motor neuron firing has been shown to lead to capture of only retrogradely-moving DCVs (Shakiryanova et al., 2006). Perhaps in *klc-1(-)* there is a loss of a signal that would normally promote dynein pausing at synapses. Finally, it is possible that KLC-1 in some way influences the balance of activity of dynein and kinesin-1, perhaps via a shared cargo adaptor, as has been recently reported for troponin-C regulation of *oskar* mRNA transport in *Drosophila* (Heber et al., 2024). For membrane transport, obvious candidate adaptors would be JIPs (Arimoto et al., 2011; Barkus et al., 2008; Celestino et al., 2022; Edwards et al., 2015a; Edwards et al., 2013; Fu and Holzbaur, 2013; Sure et al., 2018). Whether worm KLC-1 influences any of these components or processes is an important question for future research.

### Effects of kinesin-1 mutants and IDA-1::GFP expression on lifespan

The insulin/IGF-1 signalling pathway controls lifespan in many species including *C. elegans* (Kenyon, 2010b; Miller et al., 2020). The key components are the insulin-like peptides (ILPs), such as INS-6 and -7 (Artan et al., 2016; Borbolis et al., 2020; Chen et al., 2013; Pandey et al., 2024), which bind to the DAF-2 insulin/IGF-1 receptor, down-regulating the activity of the transcription factor DAF-16 and shortening life (Artan et al., 2016; Borbolis et al., 2020; Chen et al., 2013; Kenyon, 2010b; Miller et al., 2020). There is a cell autonomous feedback loop, since active DAF-16 down-regulates *ins-7* transcription (Murphy et al., 2003). However, the pathway also acts cell non-autonomously, with communication between neurons and the intestine being vital (Artan et al., 2016; Libina et al., 2003; Miller et al., 2020; Wolkow et al., 2000). Importantly, the ILPs are transported and secreted by DCVs.

A striking observation from our lifespan experiments was that the ∼2-3-fold overexpression of IDA-1::GFP (Zahn et al., 2004) shortened lifespan considerably (Fig. 6). Over-expression of the human IDA-1 homologue, IA-2, increased the number of DCVs, insulin content, and insulin secretion in MIN-6 pancreatic beta cells (Harashima et al., 2005). In *C. elegans*, IDA-1 expression levels correlate with the number of DCVs in the ventral cord region, being reduced by 50% in *ida-1* deletions and 2-fold upregulated in a mutant that expresses 8-fold more IDA-1 (Cai et al., 2009). Although *ida-1* loss gives only subtle phenotypic effects, it enhanced the phenotype of weak alleles of genes involved in insulin-like-signalling-based control of dauer larval formation (Cai et al., 2004), which happens under conditions of overcrowding, starvation, or elevated temperature. Since higher levels of IGF-1 signalling shorten lifespan in worms (Kenyon, 2010b), this may explain the reduced lifespan of *ida-1::gfp* animals, which express more IDA-1 protein and so may secrete more DCV cargo proteins such ILPs.

Where is IDA-1::GFP having its effect? It is not expressed in all neurons with DCVs (Cai et al., 2004), but is highly expressed in the ALA, VC, HSN and PHC neurons (Cai et al., 2004; Zahn et al., 2004): of these, the ALA is most closely linked to lifespan. The ALA neuron controls a form of sleep that worms undergo when they are moulting during larval development and following periods of stress that is induced by secretion of EGF and multiple neuropeptides (Iannacone et al., 2017; Konietzka et al., 2020; Nath et al., 2016; Nelson et al., 2014; Van Buskirk and Sternberg, 2007). Recovery from such stress is impaired without the ALA neuron (Hill et al., 2014) and it is the only neuron needed to inhibit feeding during stress-induced quiescence (Trojanowski et al., 2015). The ALA neuron is suggested to be a GABA recycling neuron (Gendrel et al., 2016), although it has few chemical synapses (Ramirez-Suarez et al., 2019). Instead it acts as a hub for neuropeptide signalling, both receiving and transmitting signals (Randi et al., 2023; Ripoll-Sanchez et al., 2023). It secretes many different neuropeptides including ILPs (Konietzka et al., 2020; Nath et al., 2016; Ripoll-Sanchez et al., 2023) which are released into the pseudocoelom, allowing the neuron to signal to multiple tissues, such as the intestine, at a distance (Ramirez-Suarez et al., 2019). It is activated by neuropeptides released from the IL1, SAA, RIA and URX neurons (Randi et al., 2023), and the RIA and URX neurons have been shown to control olfactory learning via INS-6 and -7 signalling to DAF-2 (Chen et al., 2013).

Intriguingly, crossing the short-lived *ida-1::gfp* strain with either *klc* mutant extended lifespan over wild-type and *klc* single mutants (which had normal longevity). Since anterograde DCV motility is reduced in the *klc-2(rf)* background, this could decrease delivery of DCVs and therefore secretion of IGF-1, so extending lifespan. In addition, UNC-116/KLC-2-mediated transport of a splice form of the DAF2 insulin receptor, DAF2c, from the cell body to the synaptic region of the ASER neuron, is required for taste avoidance learning, and this splice form can rescue effects of a *daf-2* mutation on lifespan (Ohno et al., 2014). However, both these effects would also be expected in the *klc-2(rf)* single mutant as well, which has normal lifespan. Notably, IDA-1::GFP expression improved the swimming performance of *klc-2(rf)* animals, demonstrating an overall improvement in fitness.

Expressing the *ida-1::gfp* transgene in the *klc-1(-)* background gave the strongest extension of lifespan. Although KLC-1 is important for mitochondrial transport (Zhao et al., 2021) which could affect longevity in *klc-1(-); ida-1::gfp* worms through altered mitochondrial dynamics (Byrne et al., 2019), *klc-1(-)* lifespan is normal. However, although *klc-1* is expressed at low levels in most neurons (Taylor et al., 2021), its expression is high in the IL1 and NSM neurons (Taylor et al., 2021) that control lifespan in the context of cold stress (Zhang et al., 2018) and in the ASJ neuron that responds to dietary restriction (Artan et al., 2016). All these neurons also express *ida-1* (Taylor et al., 2021). Whether the absence of KLC-1 affects longevity in response to cold and dietary restriction will be interesting to determine. Furthermore, KLC-1 is highly expressed in the gonad (Cao et al., 2017), and clear links exist between the gonad and lifespan (Kenyon, 2010a). Teasing apart the mechanisms by which the *ida-1::gfp* transgene and the *klc* mutants act synergistically to extend lifespan is a challenge for the future.

In contrast to the normal longevity of the *klc-1(-)* and *klc-2(rf)* strains, *unc-116(rf)* worms died significantly sooner (50% survival at 14 days, compared to 18 days for wild-type and *klc* mutants). This could be due in part to impaired mitochondrial dynamics. There is also a correlation between mobility, healthspan and longevity (Hahm et al., 2015; Oswal et al., 2022). The effect of *unc-116(rf)* is less severe than two dynein mutants, however, which had 50% survival rates of 9-10 days, compared to controls (50% survival at 17 days) (Koushika et al., 2004), but worse than the mild *unc-104(wy711)* allele (Li et al., 2016). In the latter case, expressing additional normal UNC-104 protein from the *rab-3* promoter slightly extended lifespan over wild-type (as well as improving associative learning and memory), in a pathway downstream of DAF-2/insulin-IGF-1-like receptor (Li et al., 2016).

Taken together, these results show that motor protein function, unsurprisingly perhaps, is important for normal lifespan, and that it interacts in complex ways with the overexpression of a DCV component that is crucial for neuropeptide secretion. Where, and how, this synergism occurs remains to be determined, but it emphasises the importance of DCV function and motility which we demonstrate relies on both kinesin-1 and kinesin-3 in C. elegans, as it does in other species.

## Methods

### *C. elegans* culture, maintenance and genetics

All *C. elegans* strains were maintained at standard conditions as previously described (Brenner, 1973): worms were kept at 20°C on 6 cm plates filled with nematode growth media (NGM) with OP-50 bacteria as food.

The *klc-1(ok2609)* (Consortium, 2012) and *klc-2(km11)* (Sakamoto et al., 2005) strains were obtained from the Caenorhabditis Genetics Centre (CGC). The *ida-1::gfp* originated from John Hutton’s laboratory, University of Colorado Health Sciences Center, Denver, and was kindly provided by Howard Davidson. The strain *unc-116(rh24sb79)* (Yang et al., 2005) was a gift from Frank McNally at the University of California, Davis. Crosses were generated for this study in the Poulin lab. The N2 Bristol variety was used as a wild-type reference strain. All strains used in this study are listed in Table 2.

**Table 2.**
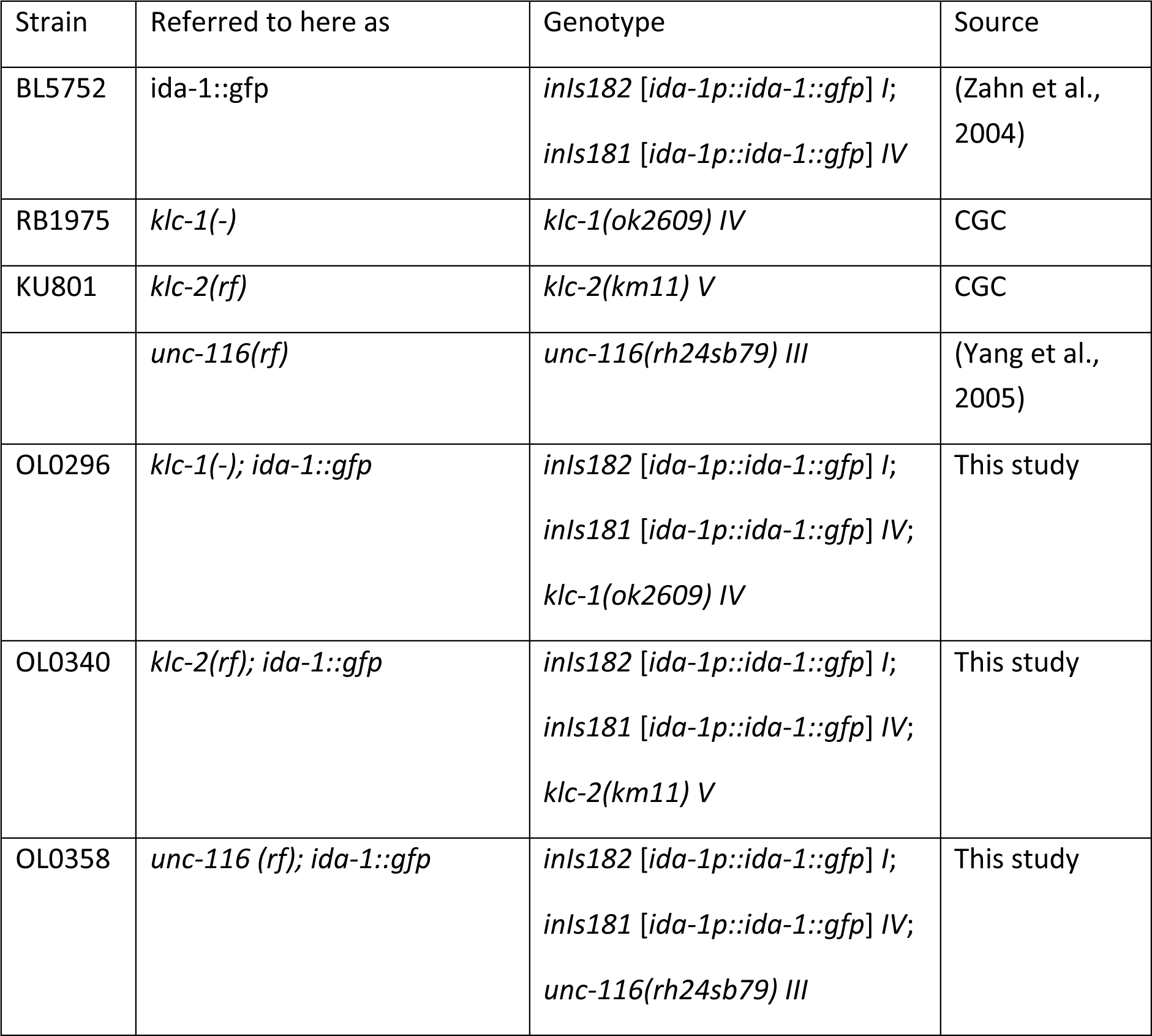
*C. elegans* strains used and generated in this study.

| Strain | Referred to here as | Genotype | Source |
| --- | --- | --- | --- |
| BL5752 | *ida-1::gfp* | *inIs182 [ida-1p::*ida-1::gfp*] I;*<br>*inIs181 [ida-1p::*ida-1::gfp*] IV* | (Zahn et al., 2004) |
| RB1975 | *klc-1(-)* | *klc-1(ok2609) IV* | CGC |
| KU801 | *klc-2(rf)* | *klc-2(km11) V* | CGC |
|  | *unc-116(rf)* | *unc-116(rh24sb79) III* | (Yang et al., 2005) |
| OL0296 | *klc-1(-); ida-1::gfp* | *inIs182 [ida-1p::*ida-1::gfp*] I;*<br>*inIs181 [ida-1p::*ida-1::gfp*] IV;*<br>*klc-1(ok2609) IV* | This study |
| OL0340 | *klc-2(rf); ida-1::gfp* | *inIs182 [ida-1p::*ida-1::gfp*] I;*<br>*inIs181 [ida-1p::*ida-1::gfp*] IV;*<br>*klc-2(km11) V* | This study |
| OL0358 | *unc-116 (rf); ida-1::gfp* | *inIs182 [ida-1p::*ida-1::gfp*] I;*<br>*inIs181 [ida-1p::*ida-1::gfp*] IV;*<br>*unc-116(rh24sb79) III* | This study |

### Worm genotyping by single worm PCR

To verify worm mutations and check successful crosses we used single worm PCR with different primers for each mutation. The *klc-1(ok2609)* null mutation is an insertion of TTTAATGTGAAA and a 1383 bp deletion. Worms with this mutation do not have any phenotype. The primers used to identify *klc-1(ok2609)* mutation are taken from the CGC: Outer Left Sequence is AAATGGCGCCAAGAGATATG, Outer Right Sequence is TCACGCAATTTTGAGTTCGT.

The *klc-2(km11)* strain exhibits a reduced-function mutation, characterized by the presence of two copies of truncated *klc-2* genes (Sakamoto et al., 2005). The endogenous *klc-2* locus carries an approximate 1.4-kilobase deletion, which truncates the protein product immediately after the coiled-coil region and eliminates the entire TPR region. Additionally, the second copy of the *klc-2* gene, inserted upstream of the endogenous locus, lacks the two alternative last exons, resulting in a protein that is truncated after the TPR domains (Sakamoto et al., 2005). These worms have a slightly uncoordinated phenotype. To follow the *klc-2(km11)* deletion, we used KLC-2-f4 (CGCCACGATCTCCTGTATTTCAATAGC) and KLC-2-r4 (GATGACGGAGTACAATGTCGAGCAAC) primers.

The reduced-function *unc-116(rh24sb79)* mutant contains two missense mutations within the motor domain from the *rh24* allele with an additional missense mutation within the motor domain (*sb79*) which suppresses the *rh24* gain-of-function allele (Yang et al., 2005). This strain serves as a hypomorphic and revertant allele. To identify this *unc-116* mutation in crosses we used its strong uncoordinated phenotype.

### Determination of lifespan

To perform lifespan experiments, 100-130 L4 worms for each strain were divided into 6 plates and were monitored daily. To verify the result, we performed three rounds of the experiment (3150 worms in total). The worms were moved to new plates if they had progeny or if the plate became contaminated. Each day the number of live, dead, and censored worms was scored. Reasons for censoring are: the worm died on the wall of the dish; the worm crawled into the agar; a bag of worms (progeny did not hatch and develop inside the mother); exploding/bursting vulva. Data were analysed using the OASIS 2 online application for survival analysis, which also performs statistical tests on the data (Han et al., 2016).

### Imaging and data analysis

Time-lapse microscopy of worms expressing IDA-1::GFP was used to capture DCV transport events. Confocal imaging was performed on a 3i inverted Spinning Disk Confocal Microscope and a motorised stage for live-cell imaging. Images were acquired using a CSU-X1 spinning disc confocal (Yokogawa) on a Zeiss Axio-Observer Z1 microscope with a 20x/0.5 EC PlanNeofluar, 40x/1.30 Plan-Apochromat (Oil; DIC), 63x/1.40 Plan-Apochromat and 100x/1.30 Plan-Neofluar objectives, Prime 95B Scientific CMOS (1200 x1200 11 µm pixels; backlit; 16-bit) camera (Photometrics) and motorised XYZ stage (ASI). The 488 nm laser was controlled using an AOTF through the laser stack (3i) allowing both rapid ‘shuttering’ of the laser and attenuation of the laser power. SlideBook software (3i) was used to capture images.

To capture images of whole worms, Multiple XY Location Capture in SlideBook was used to generate montage images (Fig. 1, S1). Montage fluorescent images (Fig. 1 A’’-D’’) were generated automatically, and montage brightfield images (Fig. 1A’-D’) were generated manually. Brightfield and fluorescence images were taken with the same camera at 100 ms exposure, with LED brightfield illumination. Brightfield images are in one focal plane, whereas fluorescence images have several z-planes combined by maximum projection.

Videos of DCV movement collected using the 100x objective were made using a single plane with 156 ms frame interval for 500 frames. Raw TIFF files were stabilized using image stabilizer plugin for ImageJ (http://www.cs.cmu.edu/~kangli/code/Image_Stabilizer.html) using the best suitable initial reference slice. For sections where stabilization did not work, for example if the worm moved significantly, the frames were deleted. Standard parameters for image stabilizer were used. A segmented line was then superimposed over the neuron in the stabilized videos and the ImageJ Multi Kymograph plugin was used to create kymographs. The best quality kymographs were chosen for subsequent analysis: 82 of *ida-1::gfp*, 66 kymographs of *klc-1(-); ida-1::gfp*, 127 kymographs of *klc-2(rf); ida-1::gfp*, and 53 kymographs of *unc-116(rf); ida-1::gfp*. Tracks were identified using KymoButler (Jakobs et al., 2019), with examples shown in Fig. 2 and Fig. S2. The KymoButler code was obtained from https://github.com/ and run in Mathematica to give the x and t positions for each track in every kymograph.

For each strain we combined KymoButler results to analyse and visualize results using custom-written Python code (the analysis code will be provided in Github upon publication). In cases where two x positions corresponded to the same t, indicating that a particle occupied two positions simultaneously, which is impossible, the average position was computed. As KymoButler results are in pixels, they were next converted to seconds and µm. In Fig. 3, all tracks for each strain were combined with one starting point to visualize transport properties and compare them across different strains.

To compare retrograde and anterograde movement, each track was divided into segments: retrograde, anterograde and stationary, using a sliding window algorithm with a 0.6444 µm movement threshold and a 2 s time threshold. Each track can have multiple segments of each type (See Fig. S3E for examples). Venn diagrams were employed to illustrate the distribution of tracks exhibiting retrograde, anterograde, stationary or combinations of segment types (Fig. S3). Velocities were computed for retrograde or anterograde segments by dividing the distance displacement by the time displacement. Tracks with fewer than four points were omitted from velocity calculations to enhance robustness (*ida1::gfp*, 37 tracks [0.5%]; *klc-1(-); ida-1::gfp*, 21 tracks [0.45%] and *klc-2(rf); ida-1::gfp*, 28 tracks [0.32%]). No tracks were excluded for *unc-116(rf); ida-1::gfp*. The velocities of retrograde or anterograde segments were averaged to derive the mean and standard deviation, along with the median velocity and maximum retrograde or anterograde velocities. The velocity distribution patterns were analysed using Gaussian Mixture Models (GMMs). The analysis was conducted separately for retrograde and anterograde movements using Python. For each strain and direction, the Akaike Information Criterion (AIC) was used to determine the optimal number of components for the GMM.

For the data plotted in Fig. S4, data were grouped according to the distance from the ALA cell body for each kymograph: <250 µm, 250-450 µm, 450-650 µm, and >650 µm. Worms of the strains used in this study have different average body lengths (n=100 worms, ±SEM): 1.12 ± 0.01 mm, *ida-1::gfp*, 1.12 ± 0.006 mm *klc-1(-); ida-1::gfp*, 0.99 ± 0.007 mm *klc-2(rf); ida-1::gfp*, and 0.77 ± 0.004 mm *unc-116(rf); ida-1::gfp*. This is why in Fig. S4 there are no kymographs for *unc-116(rf); ida-1::gfp* further than 650 µm from the ALA cell body.

Swimming or thrashing assays were performed using a Leica M165 FC Fluorescent Stereo Microscope, optiMOS™ Scientific CMOS Camera and PLANAPO 1.0x objective. Image streams were captured using Micro-Manager, consisting of 1000 frames with 15-17 ms exposure. The videos were analyzed using the Stack Deflicker and wrMTrck plugins in Fiji to quantify body bends per second for each worm. Subsequently, the data from multiple videos was combined for each strain.

## Statistical analysis

Velocity distributions of DCVs in the different strains were compared by the pairwise two-sample Kolmogorov-Smirnov (KS) test (Table S1). The null hypothesis for the KS test (the two distributions come from the same population) was rejected at the P=0.05 level. To test the significant differences in the body bends per second for each pair of strains in the swimming experiment (Fig. 5), an independent two-sample t-test was employed with the null hypothesis of no difference in the population means at the significance level of 0.05 (Table. S2). Lifespan survival functions were compared by the Log Rank Test (Table S3) with the null hypothesis of no difference in survival between two groups at the significance level of 0.05 using the online application for survival analysis, OASIS 2 (Han et al., 2016).

## Acknowledgements

We are grateful to Howard Davidson (University of Colorado Health Sciences Center, Denver), Frank McNally (University of California, Davis) and the CGC for providing worm strains. The CGC is funded by the NIH Office of Research Infrastructure Programs (P40 OD010440). VA would like to thank Dan Starr and Jon Scholey (University of California, Davis) for introducing her to imaging DCVs in *C. elegans*. The 3i spinning disk microscope used in this study was purchased by the University of Manchester Strategic Fund. Special thanks go to Dr Peter March for his help with the confocal imaging. We are very grateful to Dmytro Chekunov for help with Python and Mathematica coding.

## Competing interests

No competing interests declared.

## Funding

This work was funded by a Wellcome Trust Ph.D. studentship to AG (Grant No. 108867/Z/15/Z), and a Biotechnology and Biological Sciences Research Council CASE award, co-funded by 3i, to AB. VA acknowledges funding from a Wellcome Trust ISSF award.

## Data availability

Source data will be made available upon publication.

## Notes

### Competing Interest Statement

The authors have declared no competing interest.

### Summary of Updates

This revision now includes the supplemental material file. The main article pdf has not changed.

