## Supplemental figures 1-5 and supplementary tables 1-3 for "The role of Kinesin-1 in neuronal dense core vesicle transport and lifespan regulation in *C. elegans*"

**Table S1. Statistical analysis of DCV velocities**. P-values from pairwise Kolmogorov-Smirnov tests for velocity distribution of DCVs in the different strains. Grey colour represents insignificant differences in velocities.

| Strains | Retrograde | Anterograde |
|---|---|---|
| *ida-1::gfp* vs *klc-1(-); ida-1::gfp* | 4.59e-14 | 0.06 |
| *ida-1::gfp* vs *klc-2(rf); ida-1::gfp* | 4.81e-63 | 4.82e-15 |
| *ida-1::gfp* vs *unc-116(rf); ida-1::gfp* | 1.14e-248 | 2.94e-101 |
| *klc-1(-); ida-1::gfp* vs *klc-2(rf); ida-1::gfp* | 3.8e-63 | 6.82e-21 |

**Table S2. Statistical analysis of worm swimming**. P-values from an independent two-sample t-test to show significant differences in the body bends per second for each pair of strains (shown in Fig. 5). Number of worms: N2, 49 worms; *ida-1::gfp,* 49 worms; *klc-1(-),* 53 worms; *klc-2(rf),* 30 worms; *unc-116(rf),* 38 worms; *klc-1(-); ida-1::gfp,* 51 worms; *klc-2(rf); ida-1::gfp,* 49 worms and *unc-116(rf); ida-1::gfp,* 43 worms. Non-significant comparisons are shown with grey shading.

| Stain | N2 | *ida-1:: gfp* | *klc-1(-)* | *klc-1(-); ida-1:: gfp* | *klc-2(rf)* | *klc-2(rf); ida-1:: gfp* | *unc116(rf)* | *unc-116(rf); ida-1::gfp* |
|---|---|---|---|---|---|---|---|---|
| N2 | | 0.728 | 0.872 | 0.287 | 1.14e-21 | 4.42e-15 | 9.35e-46 | 9.97e-45 |
| *ida-1::gfp* | 0.728 | | 0.671 | 0.206 | 1.35e-21 | 4.69e-14 | 6.87e-42 | 2.92e-41 |
| *klc-1(-)* | 0.872 | 0.671 | | 0.424 | 1.84e-21 | 4.18e-13 | 1.66e-37 | 2.72e-37 |
| *klc-1(-); ida-1::gfp* | 0.287 | 0.206 | 0.424 | | 3.17e-21 | 2.44e-13 | 7.55e-34 | 9.62e-34 |
| *klc-2(rf)* | 1.14e-21 | 1.35e-21 | 1.84e-21 | 3.17e-21 | | 3.22e-05 | 7.01e-16 | 6.78e-16 |
| *klc-2(rf); ida-1::gfp* | 4.42e-15 | 4.69e-14 | 4.18e-13 | 2.44e-13 | 3.22e-05 | | 1.55e-25 | 1.57e-25 |
| *unc-116(rf)* | 9.35e-46 | 6.87e-42 | 1.66e-37 | 7.55e-34 | 7.01e-16 | 1.55e-25 | | 0.116 |
| *unc-116(rf); ida-1::gfp* | 9.97e-45 | 2.92e-41 | 2.72e-37 | 9.62e-34 | 6.78e-16 | 1.57e-25 | 0.116 | |

**Table S3. Statistical analysis of worm lifespan data**. P-values from a Log-Rank test between the survival functions for each pair of strains from Fig. 6. The null hypothesis was that there was no difference in survival functions. For the comparisons indicated as <α, this corresponds to a P-value of < 1.0 x $10^{-10}$. Non-significant comparisons are shown with grey shading.

| Strain | N2 | *ida-1::gfp* | *klc-1(-)* | *klc-2(rf)* | *unc-116(rf)* | *klc-1(-); ida-1:: gfp* | *klc-2(rf); ida-1:: gfp* | *unc-116(rf); ida-1::gfp* |
|---|---|---|---|---|---|---|---|---|
| N2 | | 0.0003 | 0.4971 | 0.8112 | <α | <α | 1.2e-6 | <α |
| *ida-1::gfp* | 0.0003 | | 3.3e-5 | 0.0002 | <α | <α | <α | <α |
| *klc-1(-)* | 0.4971 | 3.3e-5 | | 0.5729 | <α | <α | 3.2e-5 | <α |
| *klc-2(rf)* | 0.8112 | 0.0002 | 0.5729 | | <α | <α | 4.6e-6 | <α |
| *unc-116(rf)* | <α | <α | <α | <α | | <α | <α | 0.0286 |
| *klc-1(-); ida-1::gfp* | <α | <α | <α | <α | <α | | 0.0202 | <α |
| *klc-2(rf); ida-1::gfp* | 1.2e-6 | <α | 3.2e-5 | 4.6e-6 | <α | 0.0202 | | <α |
| *unc-116(rf); ida-1::gfp* | <α | <α | <α | <α | 0.0286 | <α | <α | |

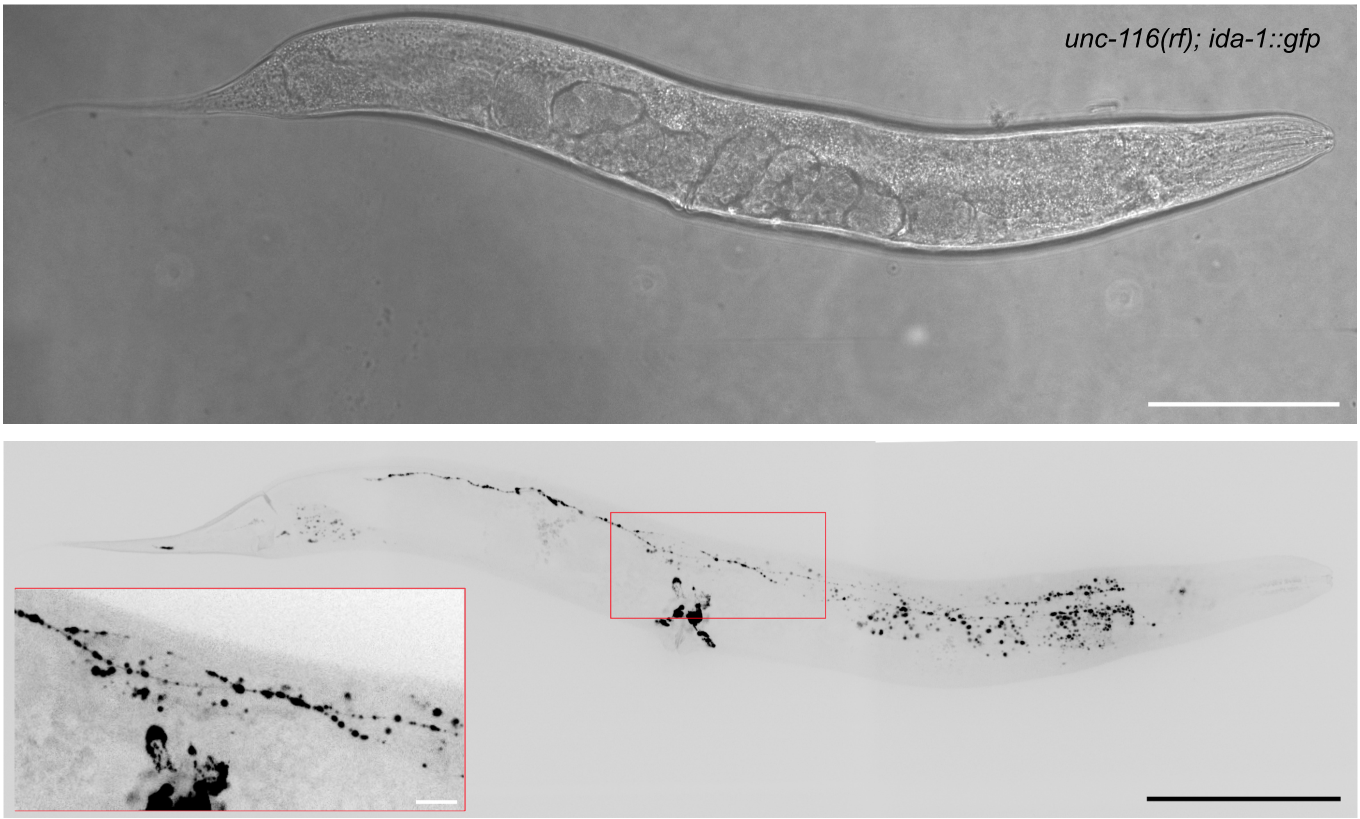


**Figure S1. Additional brightfield and fluorescence images of the *unc-116(rf); ida-1::gfp* strain.** An enlarged section of each ALA neuron is shown in the red box. The lighter spots in the brightfield image are artefacts generated by the tiling of multiple images. The larger particles visible outside the ALA neuron are auto-fluorescent granules, mainly located in the gut. Scale bars are 100 µm with a 10 µm scale bar in the inset.

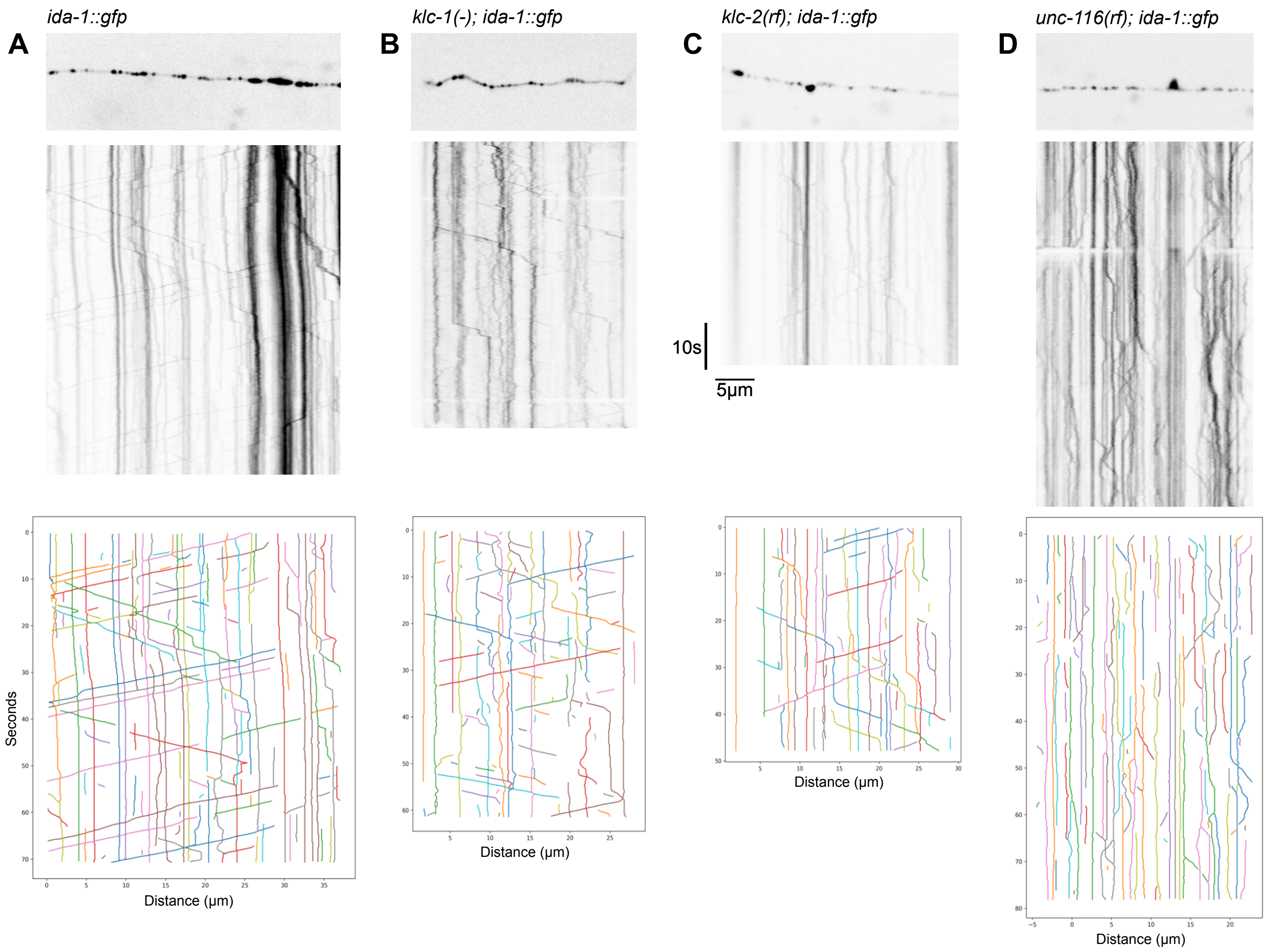


**Figure S2. Additional kymographs of DCV movement in the ALA neuron in different strains**. The initial frame of each movie is shown on top, with the kymograph below. A) *ida-1::gfp*, B) *klc-1(-); ida-1::gfp*, C) *klc-2(rf); ida-1::gfp*, and D) *unc-116(rf); ida-1::gfp*. The nerve terminal (microtubule plus ends) is on the right for each kymograph. Kinesin-driven motile DCVs correspond to lines sloping from top left to bottom right, with dynein-driven lines sloping from top right to bottom left. Vertical lines indicate stationary DCVs (individual or clusters). Scale bars are 5 μm.

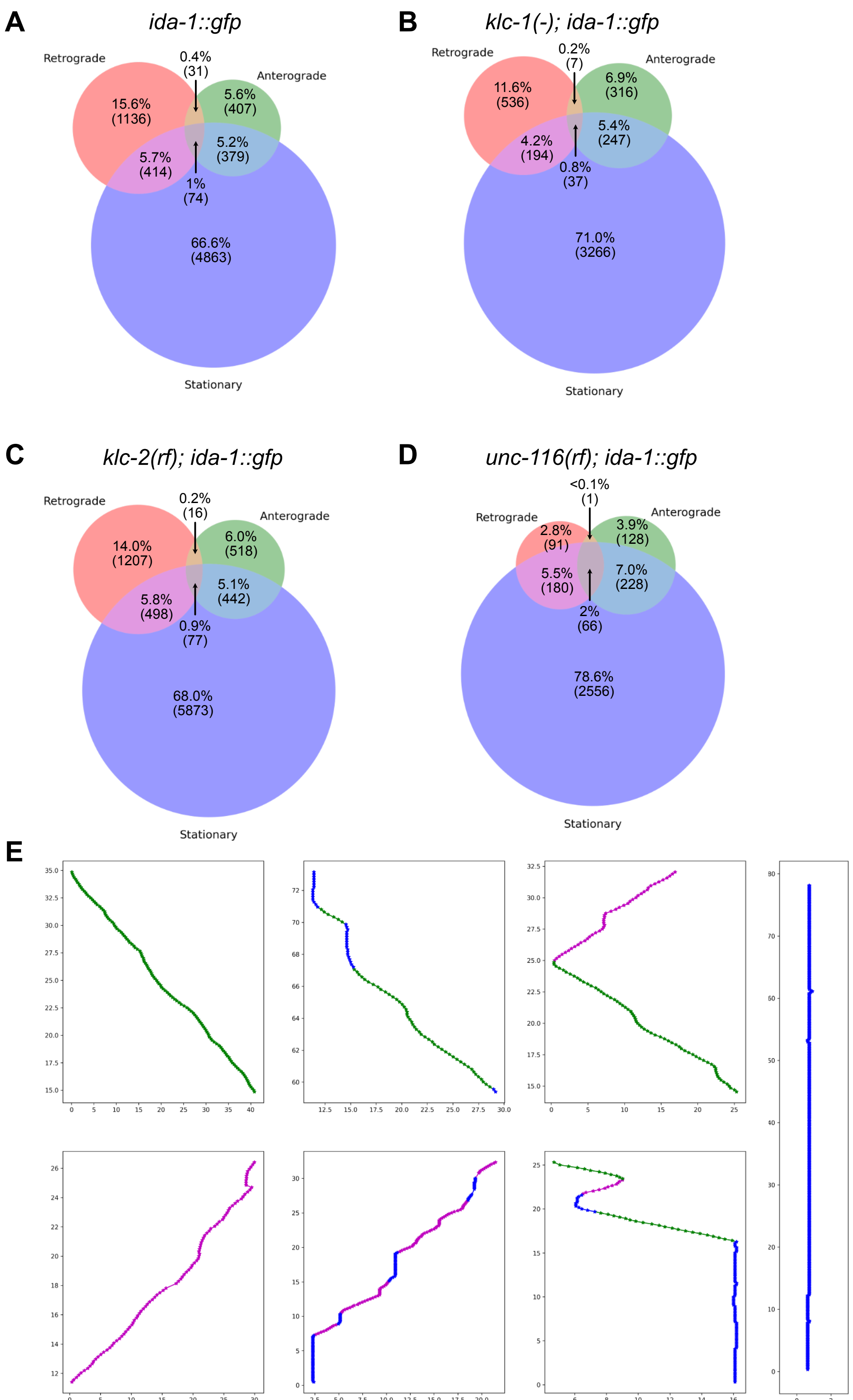


**Figure S3. Analysis of DCV motility.** The proportion of tracks with retrograde, anterograde and stationary segments for each of the four strains, displayed as Venn diagrams. A) *ida-1::gfp*, B) *klc-1(-); ida-1::gfp*, C) *klc-2(rf); ida-1::gfp*, and D) *unc-116(rf); ida-1::gfp*. E) Example *ida-1::gfp* tracks showing different types of movement: retrograde segments are shown in green, anterograde in magenta and stationary segments are in blue. Seven types of tracks include: only retrograde movement; retrograde and stationary movement; retrograde and anterograde movement; only anterograde movement; anterograde and stationary movement; retrograde, anterograde and stationary movement; only stationary movement. Each track can have multiple segments of each type.

### Figure S4

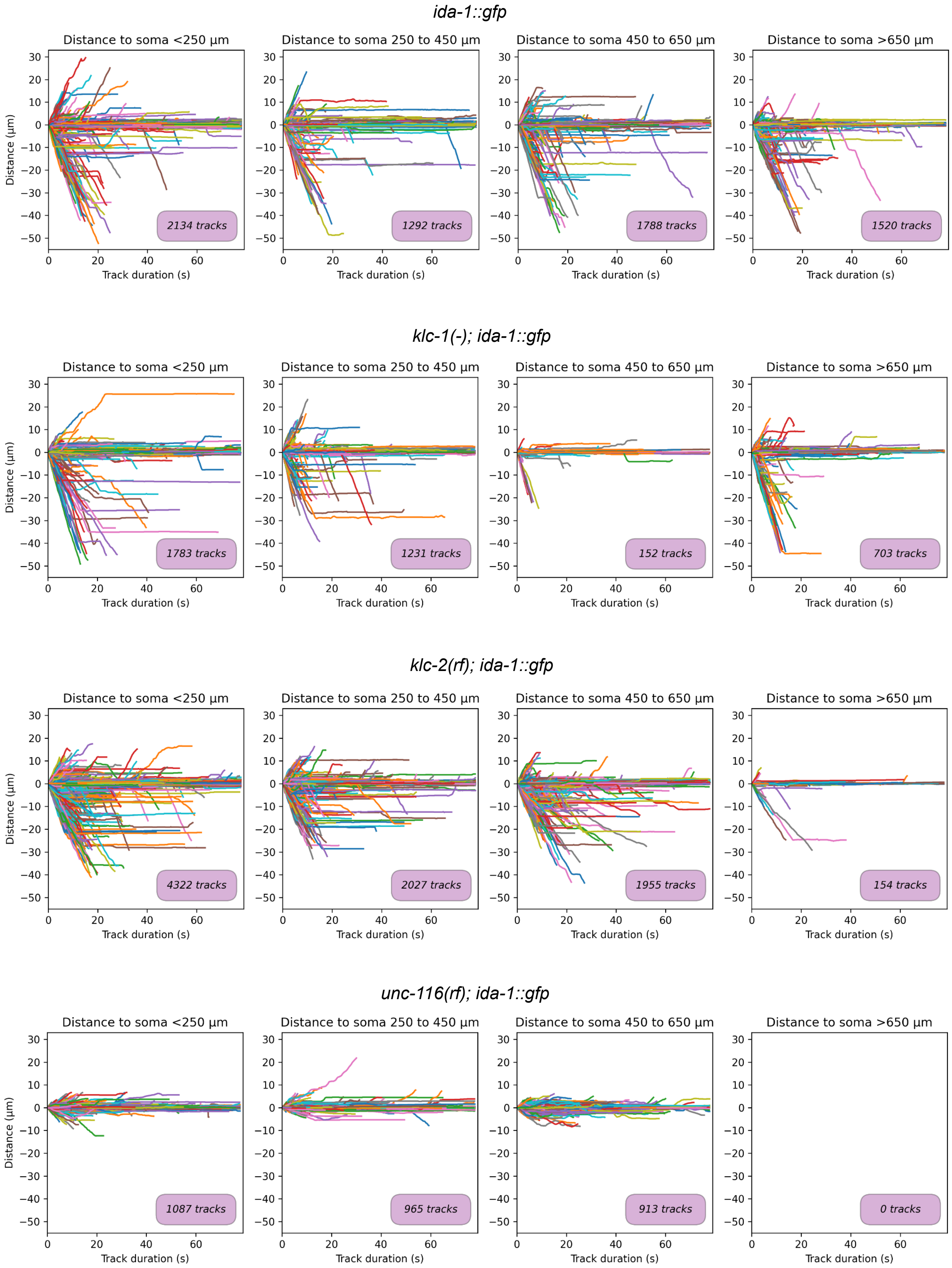
ida-1::gfp
Distance to soma <250 µm
Distance to soma 250 to 450 µm
Distance to soma 450 to 650 µm
Distance to soma >650 µm
2134 tracks
1292 tracks
1788 tracks
1520 tracks
klc-1(-); ida-1::gfp
Distance to soma <250 µm
Distance to soma 250 to 450 µm
Distance to soma 450 to 650 µm
Distance to soma >650 µm
1783 tracks
1231 tracks
152 tracks
703 tracks
klc-2(rf); ida-1::gfp
Distance to soma <250 µm
Distance to soma 250 to 450 µm
Distance to soma 450 to 650 µm
Distance to soma >650 µm
4322 tracks
2027 tracks
1955 tracks
154 tracks
unc-116(rf); ida-1::gfp
Distance to soma <250 µm
Distance to soma 250 to 450 µm
Distance to soma 450 to 650 µm
Distance to soma >650 µm
1087 tracks
965 tracks
913 tracks
0 tracks
Distance (µm)
Track duration (s)

### Figure S5

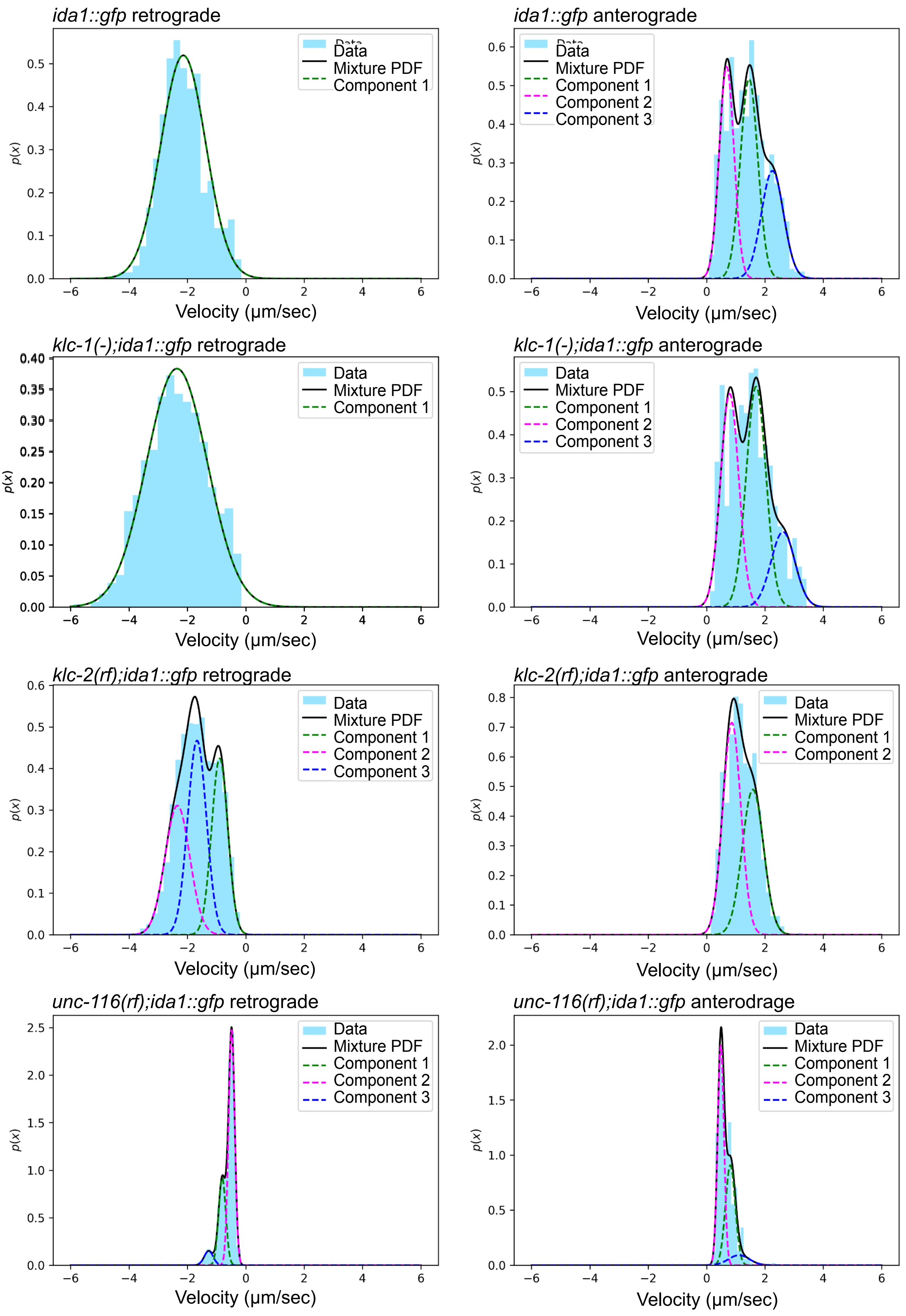

ida1::gfp retrograde
ida1::gfp anterograde
klc-1(-);ida1::gfp retrograde
klc-1(-);ida1::gfp anterograde
klc-2(rf);ida1::gfp retrograde
klc-2(rf);ida1::gfp anterograde
unc-116(rf);ida1::gfp retrograde
unc-116(rf);ida1::gfp anterodrage
Data
Mixture PDF
Component 1
Component 2
Component 3
p(x)
Velocity (μm/sec)

**Supplementary movie legends**

**Supplementary movie 1. Wild-type DCV movement**. A spinning disk confocal video of DCV movement in the ALA neuron in an *ida-1::gfp* worm. The nerve terminal (microtubule plus ends) is on the right. Kinesin-driven DCVs correspond to particles moving to the right, with dynein-driven DCVs moving to the left. The frame rate is 2 times real time. Scale bar = 5 µm.

**Supplementary movie 2. DCV movement in the *klc-1(-) mutant***. A spinning disk confocal video of DCV movement in the ALA neuron in a *klc-1(-); ida-1::gfp* worm. The nerve terminal (microtubule plus ends) is on the right. Kinesin-driven DCVs correspond to particles moving to the right, with dynein-driven DCVs moving to the left. The frame rate is 2 times real time. Scale bar = 5 µm.

**Supplementary movie 3. DCV movement in the *klc-2(rf) mutant***. A spinning disk confocal video of DCV movement in the ALA neuron in a *klc-2(rf); ida-1::gfp* worm. The nerve terminal (microtubule plus ends) is on the right. Kinesin-driven DCVs correspond to particles moving to the right, with dynein-driven DCVs moving to the left. The frame rate is 2 times real time. Scale bar = 5 µm.

**Supplementary movie 4. DCV movement in the *unc-116(rf) mutant***. A spinning disk confocal video of DCV movement in the ALA neuron in a *unc-116(rf); ida-1::gfp* worm. The nerve terminal (microtubule plus ends) is on the right. Kinesin-driven DCVs correspond to particles moving to the right, with dynein-driven DCVs moving to the left. The frame rate is 2 times real time. Scale bar = 5 µm.
